# A potent bispecific nanobody protects hACE2 mice against SARS-CoV-2 infection via intranasal administration

**DOI:** 10.1101/2021.02.08.429275

**Authors:** Xilin Wu, Lin Cheng, Ming Fu, Bilian Huang, Linjing Zhu, Shijie Xu, Haixia Shi, Doudou Zhang, Huanyun Yuan, Waqas Nawaz, Ping Yang, Qinxue Hu, Yalan Liu, Zhiwei Wu

## Abstract

The dramatically expanding COVID-19 needs multiple effective countermeasures. Neutralizing antibodies are a potential therapeutic strategy for treating COVID-19. A number of neutralizing nanobodies (Nbs) were reported for their *in vitro* activities. However, *in vivo* protection of these nanobodies was not reported in animal models. In the current report, we characterized several RBD-specific Nbs isolated from a screen of an Nb library derived from an alpaca immunized with SARS-CoV-2 spike glycoprotein (S); among them, three Nbs exhibited picomolar potency against SARS-CoV-2 live virus, pseudotyped viruses, and 15 circulating SARS-CoV-2 variants. To improve the efficacy, various configurations of Nbs were engineered. Nb_15_-Nb_H_-Nb_15_, a novel trimer constituted of three Nbs, was constructed to be bispecific for human serum albumin (HSA) and RBD of SARS-CoV-2. Nb_15_-Nb_H_-Nb_15_ exhibited sub-ng/ml neutralization potency against the wild-type and currently circulating variants of SARS-CoV-2 with a long half-life *in vivo*. In addition, we showed that intranasal administration of Nb_15_-Nb_H_-Nb_15_ provided 100% protection for both prophylactic and therapeutic purposes against SARS-CoV-2 infection in transgenic hACE2 mice. Nb_15_-Nb_H_-Nb_15_ is a potential candidate for both prevention and treatment of SARS-CoV-2 through respiratory administration.

**One sentence summary:** Nb_15_-Nb_H_-Nb_15_, with a novel heterotrimeric bispecific configuration, exhibited potent and broad neutralization potency against SARS-CoV-2 *in vitro* and provided *in vivo* protection against SARS-CoV-2 infection in hACE2 transgenic mice via intranasal delivery.

Graphical abstract:

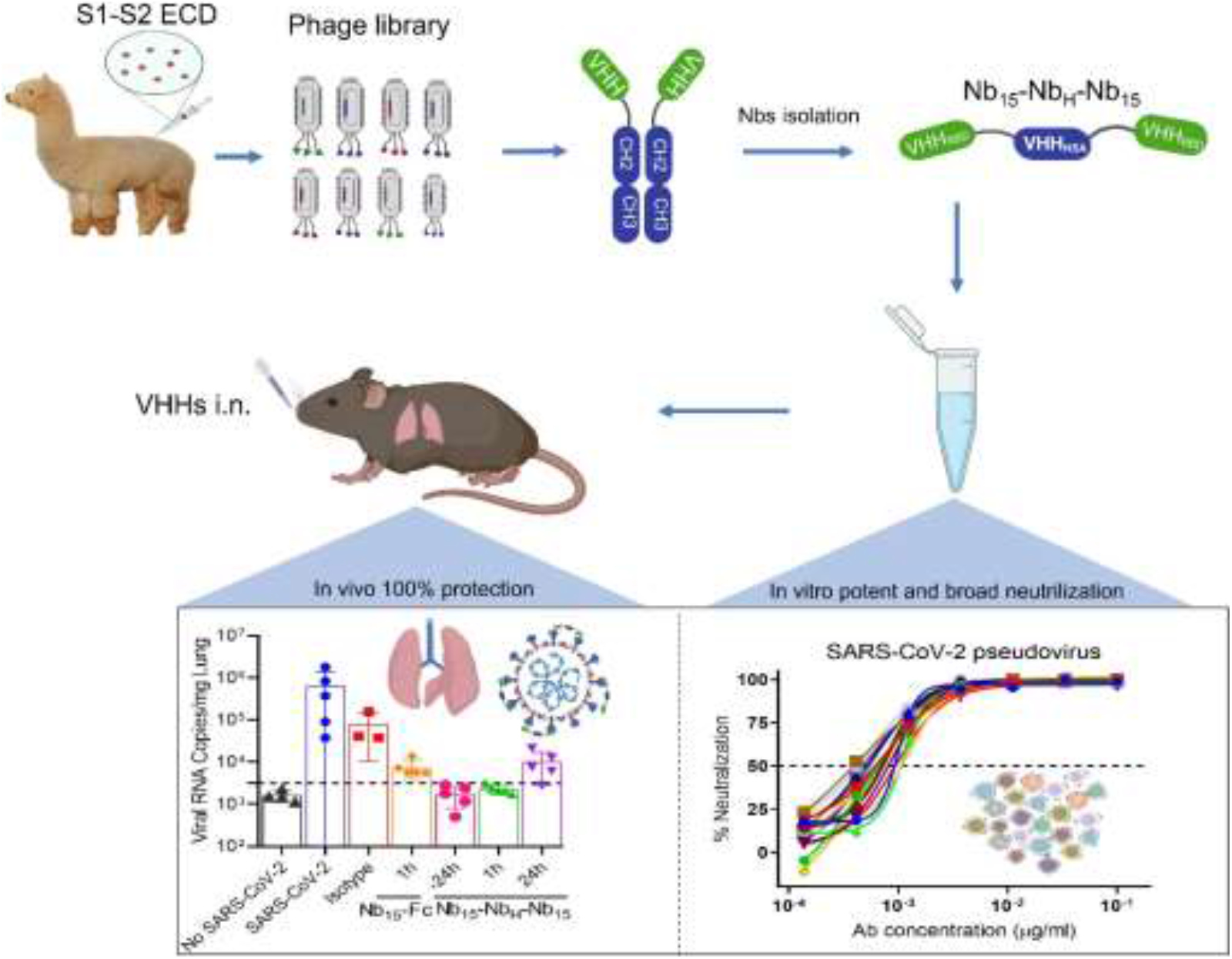

**Highlights:**

1. We described a novel heterotrimeric configuration of Nb-Nb_H_-Nb (Nb_15_-Nb_H_-Nb_15_) that exhibited improved viral inhibition and stability.
2. Nb_15_-Nb_H_-Nb_15_ provides ultrahigh neutralization potency against SARS-CoV-2 wild type and 18 mutant variants, including the current circulating variants of D614G and N501Y predominantly in the UK and South Africa.
3. It is the first to demonstrate the Nbs efficacy in preventing and treating SARS-CoV-2 infection in hACE2 transgenic mice via intranasal delivery.

## Introduction

As of Feb. 1^st^, 2021, the novel coronavirus SARS-CoV-2 has caused more than 100 million confirmed cases and over 2.2 million deaths globally. The containment of the expanding COVID-19 pandemic needs multiple countermeasures. Prophylactic vaccines have been recently approved^1^, and a number of SARS-CoV-2-neutralizing monoclonal antibodies (mAbs) that target the receptor binding domain (RBD) of spike protein^2–6^ were identified, which could be developed as either therapeutic or prophylactic agents.

In addition to conventional antibodies, camelids also generate heavy-chain-only antibodies (HCAbs), constituting a single variable domain (Nb) specific for binding antigens^7^. This single variable domain, referred as a single-domain antibody, VHH, or Nanobody (Nb), has higher affinity, thermal stability and chemostability than most antibodies^8,9^ and can easily be constructed into multivalent formats devoid of Fc, which will overcome potential deleterious antibody-dependent enhancement (ADE) of infection observed in some viral infections, including Dengue virus, HIV and SARS-CoV^10–12^. Their favorable biophysical properties have led to the development of several Nbs as therapeutics against viral infection, such as severe fever with thrombocytopenia syndrome virus (SFTSV)^13^ and respiratory syncytial virus (RSV)^14–16^.

SARS-CoV-2 is transmitted via the upper respiratory tract^17^ and analysis of clinical specimens showed that SARS-CoV-2 was detected with the highest viral copies in multiple sites of the respiratory tract while few viral copies in the blood^18^, indicating that biotherapeutic agents directly delivered via respiratory route to the sites of infection would be an attractive alternative to systemic routes of administration. Parenteral inoculation of a biotherapeutic antibody is a particularly ineffective way to deliver drugs to the respiratory tract. Indeed, a study on Mepolizumab (anti-interleukin-5 mAb) demonstrated that only 0.2% of the dose administered reached the lung via systemic administration^19^. In addition, the therapeutic effect of pulmonary delivery of human immunoglobulins for controlling RSV in cotton rats was shown to be 160 times more effective than that by the parenteral administration^20^. Thus, pulmonary delivery may be superior to parenteral administration for the treatment of respiratory tract infections. A key requirement for pulmonary delivery is the stability of the biologic to endure the degrading environment and thus the drug will have to be formulated to maintain its structural integrity and bioactivity through upper respiratory tract and the lungs. Nbs are delivered directly to the lungs via an inhaler given their small size, simple and robust structure, high thermal stability, and solubility. For instance, ALX-0171, a homotrimeric Nbs, is highly effective in reducing nasal and lung RSV titers via the pulmonary administration of inhalation^14,16^.

To date, several Nbs against SARS-CoV-2 were reported for their *in vitro* activities, but none of them have been evaluated via intranasal administration in animal models^21–26^. In the current report, anti-sera specific for RBD were elicited by immunizing an alpaca with SARS-CoV-2 spike glycoprotein (S). Nbs specific for RBD were isolated from a phage library displaying Nbs. We identified three Nbs exhibiting potent neutralization activity against live virus and a panel of SARS-CoV-2 pseudotyped viruses. To improve efficacy and stability, various configurations of Nbs were engineered. Nb_15_-Nb_H_-Nb_15_, a novel bispecific format constituted of three Nbs, was constructed to be tri-valent and bispecific for RBD of SARS-CoV-2 and human serum albumin (HSA). This novel bispecific antibody exhibited potent inhibitory activity against the wild-type and variants of SARS-CoV-2, including the currently circulating variants, such as the predominant mutant viruses in the UK and South Africa with N501Y mutation. In addition, we showed that intranasal administration of Nb_15_-Nb_H_-Nb_15_ provided 100% protection in both the prevention and treatment of SARS-CoV-2 infected transgenic hACE2 mice. Nb_15_-Nb_H_-Nb_15_ is a potential candidate for both prevention and treatment of SARS-CoV-2.

## Results

### Anti-sera response elicited by S protein

One alpaca was immunized with SARS-CoV-2 recombinant S protein (Fig. S1A). Compared to the pre-immunized serum (blank serum), the anti-serum after the third immunization exhibited specific serologic activities against SARS-CoV-2 S and RBD proteins with binding titers of 2.19 × 10^6^ and 7.29 × 10^5^, respectively (Fig. S1A and S1B). The immunized serum showed potent neutralization activity against the pseudotyped SARS-CoV-2 with a half-maximal neutralization dilution (ND_50_) of ~9600 (Fig. S1C), orders of magnitude higher than those of convalescent COVID-19 patients^27^. These data indicate that potent anti-serum specific for RBD with robust neutralization against SARS-CoV-2 was induced in the immunized alpaca.

### Isolation of Nbs with potent neutralization activity against SARS-CoV-2

To isolate monoclonal Nbs, C9-Nb library, a phage library displaying Nbs from the immunized alpaca, was constructed with a size of 2.0 × 10^9^, 100% sequence diversity, and 96% in-frame rate as validated by PCR and sequencing (Fig. S2A). Nbs specific for SARS-CoV-2 S protein were isolated through 3 rounds of biopanning on the C9-Nb phage library by S protein. The panned library was analyzed by phage ELISA for binding with S protein, and the incremental increase of the OD_450_ readout from 0.79 before enrichment to 1.6, 2.4, and 2.8 after the first, second, and third rounds of enrichment, respectively (Fig. S2B), indicating successful enrichment. To verify whether the enriched library contained specific S-reactive phages, 40 and 46 clones were selected from the libraries after the second and third rounds of enrichment for single-phage ELISA, respectively. The percentage of positive clones was 57.5% and 69.6% for the second and third rounds, respectively (Fig. S2C). Among these positive binders, 21 unique Nb sequences were identified according to the sequencing results (Table S1). To further characterize, these 21 Nbs were expressed in mammalian cells by fusing the Nb gene with a human Fc1, which was cloned into the pCDNA3.4 vector to express Nb-Fc antibody (named as Nb-Fc) (Fig. S3A). ELISA results showed that all 21 Nb-Fcs reacted with S protein; among them, 14 Nb-Fcs displayed specific binding with SARS-CoV-2 RBD protein (Fig. S3B). These results were validated by bio-layer interferometry (BLI), wherein 14 Nb-Fcs exhibited specific binding to RBD with *K*_*D*_ values ranging from 37.6 to 4.25 nM (Fig. S3C and S3D). Neutralization analysis showed potent inhibition of pseudotyped SARS-CoV-2 by culture supernatants of RBD-specific Nb_15_-Fc, Nb_22_-Fc and Nb_31_-Fc (Fig. S3E).

### Epitope analysis of Nb-Fcs

The purified Nb_15_-Fc, Nb_22_-Fc and Nb_31_-Fc exhibited dose dependent binding with RBD protein on ELISA (Fig. S4A). In addition, Nb_15_-Fc, Nb_22_-Fc and Nb_31_-Fc likely reacted with conformational structure as their bindings with reduced RBD protein were almost completely abolished (Fig. S4B). The kinetic binding of Nb_15_-Fc, Nb_22_-Fc and Nb_31_-Fc with RBD protein ranged from *K*_*D*_ of 1.13 to 1.76 nM, indicating tightly clustered binding characteristics (Fig. 4C-E), which was substantiated by the superimposed ELISA binding curves (Fig. S4A). These three Nb-Fcs were next evaluated for epitope specificity in a competition assay by BLI using RBD protein as a capture antigen. The results revealed that the pre-bound Nb-Fcs efficiently blocked the further binding of the other two Nb-Fcs to RBD protein, suggesting that all three Nb-Fcs likely recognize an overlapping epitope (Fig. S5). Together, Nb_15_-Fc, Nb_22_-Fc and Nb_31_-Fc recognize a quaternary and overlapping epitope on RBD with nanomolar affinities.

### Nb-Fcs exhibiting potent and broad neutralization against SARS-CoV-2 and variants

The neutralizing activity of Nb_15_-Fc, Nb_22_-Fc and Nb_31_-Fc against SARS-CoV-2 live virus was investigated in Vero E6 cells. All three Nb-Fcs exhibited potent neutralization activity with IC_50_ values in the range of 0.0033-0.0068 μg/ml (41.3-75 pM) and IC_90_ of 0.0156-0.0235μg/ml (195-293.8 pM) (Fig. 1A, Fig. S6 and Table S2). The neutralizing potency was validated by SARS-CoV-2 pseudovirus neutralization assay with consistent results, with IC_50_ values of 0.0008, 0.0018 and 0.0023 μg/ml (10, 22.5 and 28.8 pM), respectively (Fig. 1B and Table S2). The IC_50_ values are comparable to those of the most potent neutralizing antibodies or Nbs reported^4,21,22,27,28^. The cross neutralization of these Nbs against other coronaviruses was also evaluated in pseudovirus assay, and the results showed that these three Nb-Fcs did not inhibit either MERS-CoV or SARS-CoV pseudovirus (Fig. 1C and 1D) but inhibited 15 representative variants of SARS-CoV-2 that are identified to represent over 7000 unique viral genomes^2^. In addition, Nb_15_-Fc, Nb_22_-Fc and Nb_31_-Fc also inhibited the replication of recently arising SARS-CoV-2 variants with D614G mutation with similar potency (Fig. 1E-G and Table S2). These evidences demonstrate the broadly neutralizing activity of the Nb-Fcs against SARS-CoV-2 and suggest that the Nb-Fcs target at a highly conserved epitope on RBD protein. Taken together, all these three Nb-Fcs exhibited excellent neutralization potency against the original and the representative variants of SARS-CoV-2 while did not inhibit MERS-CoV and SARS-CoV infection. Given the overlapped epitope recognized by the three Nb-Fcs, Nb_15_-Fc with the highest neutralization potency was selected for further investigation.

**Figure 1.**
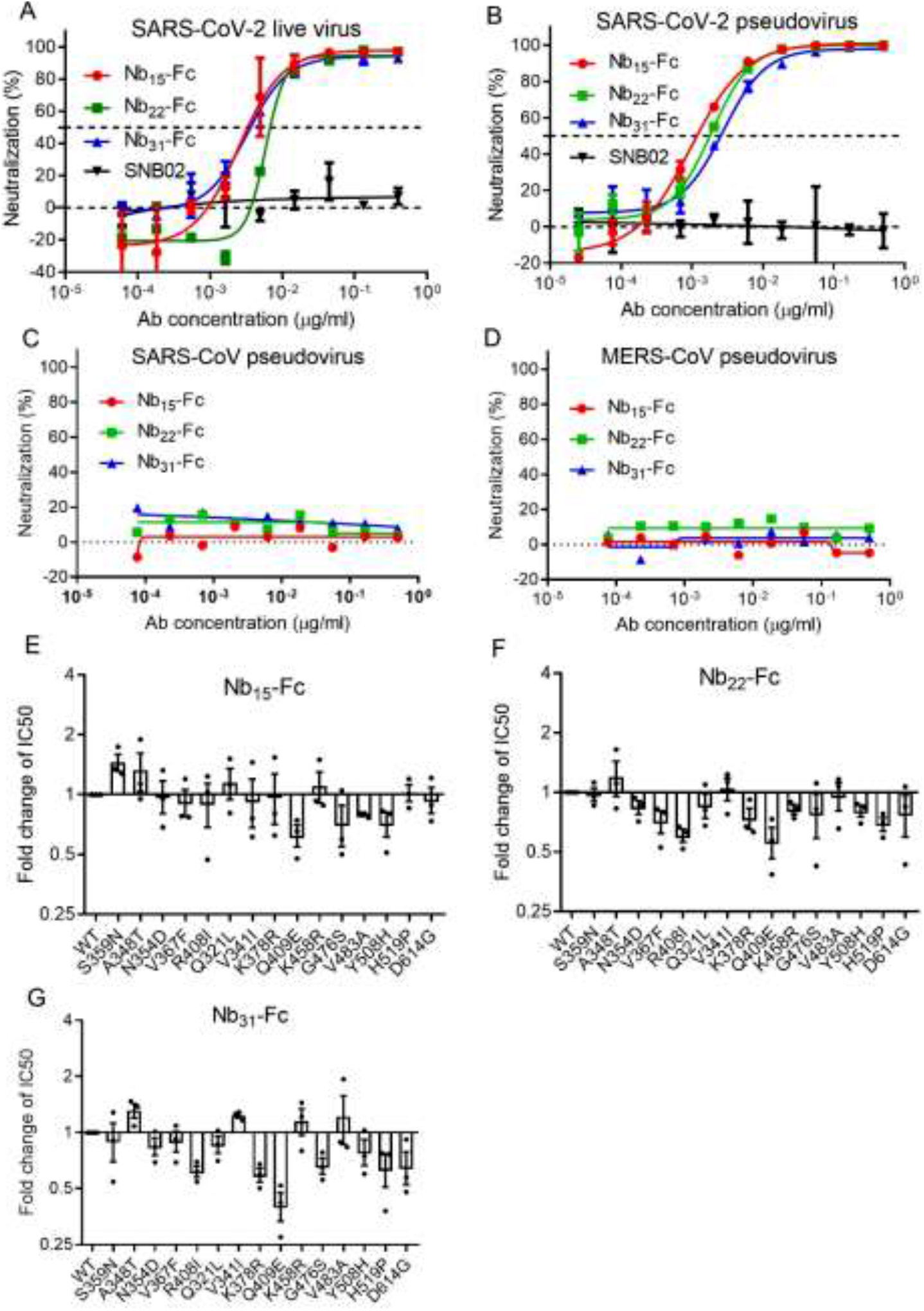
Characterizing the potency and breadth of neutralization conferred by Nb-Fcs. The neutralization potency of Nb-Fcs was detected based on authentic SARS-CoV-2 plaque reduction neutralization test (**A**) and the pseudotyped SARS-CoV-2 neutralization assay (**B**). SNB02 was taken as negative isotype control antibody (Nb fused with human Fc1). (**C**) Nb-Fcs were tested for the neutralization against the pseudovirus infection of SARS-CoV and MERS-CoV. The pseudovirus of 15 SARS-CoV-2 variants identified from circulating viral sequences were tested to evaluate the neutralization potency conferred by Nb_15_-Fc (**D**), Nb_22_-Fc (**E**) and Nb_31_-Fc (**F**), respectively. The *y* axis shows the ratio of IC_50_ of indicated SARS-CoV-2 variant/IC50 of SARS-CoV-2 wild type (WT) conferred by Nb-Fcs, The name of SARS-CoV-2 variants with amino acid point mutation based on wild type of SARS-CoV-2 were indicated. Data represent as mean ± SEM. All experiments were repeated at least twice.

### Construction and characterization of multiple-valent Nb_15_s

To improve potency, prolong *in vivo* half-life and avoid potential Fc-mediated ADE, a number of dimeric and trimeric configurations of Nbs were engineered. Monomer (1×Nb_15_), homodimer (2×Nb_15_), homotrimer (3×Nb_15_), and homotetramer (4×Nb_15_) were constructed and analyzed by BLI. The binding of these constructs to RBD protein showed an increasing *K*_*D*_ ranging from 12 to <0.001 nM as the valence increased (Fig. S7A and S7B). Multivalent formats of Nb_15_ were evaluated for neutralization against SARS-CoV-2 infection. Monomeric 1×Nb_15_ exhibited low inhibitory activity with an IC_50_ of 307 ng/ml (2.3 nM) while the bi-, tri- and tetra-valent configurations exhibited higher neutralization potency than the monomer but comparable potency among the multimers with IC_50_ values of 2.8, 3.5 and 2.3 ng/ml (11, 9.0, 4.3 pM), respectively (Fig. S7C and Table S3), suggesting that increasing valence does not confer improved anti-viral activity. As such, 3×Nb_15_ was selected for further functional exploration.

### Nb_15_-Nb_H_-Nb_15_, heterotrimer and bi-specific for RBD and HSA, exhibiting potent neutralization against SARS-CoV-2

In order to improve efficacy and stability *in vivo*, we constructed bi-specific Nbs consisting of one Nb specific for HSA (Nb_H_) developed by our lab and one or two Nb_15_s specific for RBD with (G4S)_3_ as the linker between each Nb (Fig. 2A) and analyzed their binding and viral inhibitory activities. In addition to the heterodimeric configuration of Nb-Nb_H_ that was previously reported^29^, various new configurations of Nbs were engineered as depicted in Fig. 2A. ELISA analysis showed that all combinations containing Nb_15_ reacted with RBD protein; among them, heterotrimeric Nb_15_-Nb_15_-Nb_H_, Nb_H_-Nb_15_-Nb_15_ and Nb_15_-Nb_H_-Nb_15_ exhibited better binding with RBD protein than heterodimeric Nb_15_-Nb_H_ and Nb_H_-Nb_15_ configurations (Fig. 2A and 2B). Furthermore, Nb_15_-Nb_H_, Nb_H_-Nb_15_-Nb_15_ and Nb_15_-Nb_H_-Nb_15_ were the best HSA binders as compared to other configurations (Fig. 2A and 2C). Bi-specific Nbs in various configurations were tested for the inhibition of SARS-CoV-2 infection; among them, Nb_15_-Nb_H_-Nb_15_ exhibited the most potent neutralization of the virus with an IC_50_ of 0.4 ng/ml (9.0 pM) (Fig. 2D and Table S3). We next compared Nb_15_-Nb_H_-Nb_15_ with homotrimer Nbs (3×Nb_15_) or Nb-Fc for their binding and anti-viral activities and found that 3×Nb_15_, Nb_15_-Nb_H_-Nb_15_ and Nb_15_-Fc exhibited comparable potency with IC_50_ values of 0.4, 0.4, and 0.9 ng/ml (9.0, 9.0 and 11.3 pM), respectively (Fig. 2E and Table S3).

**Figure 2.**
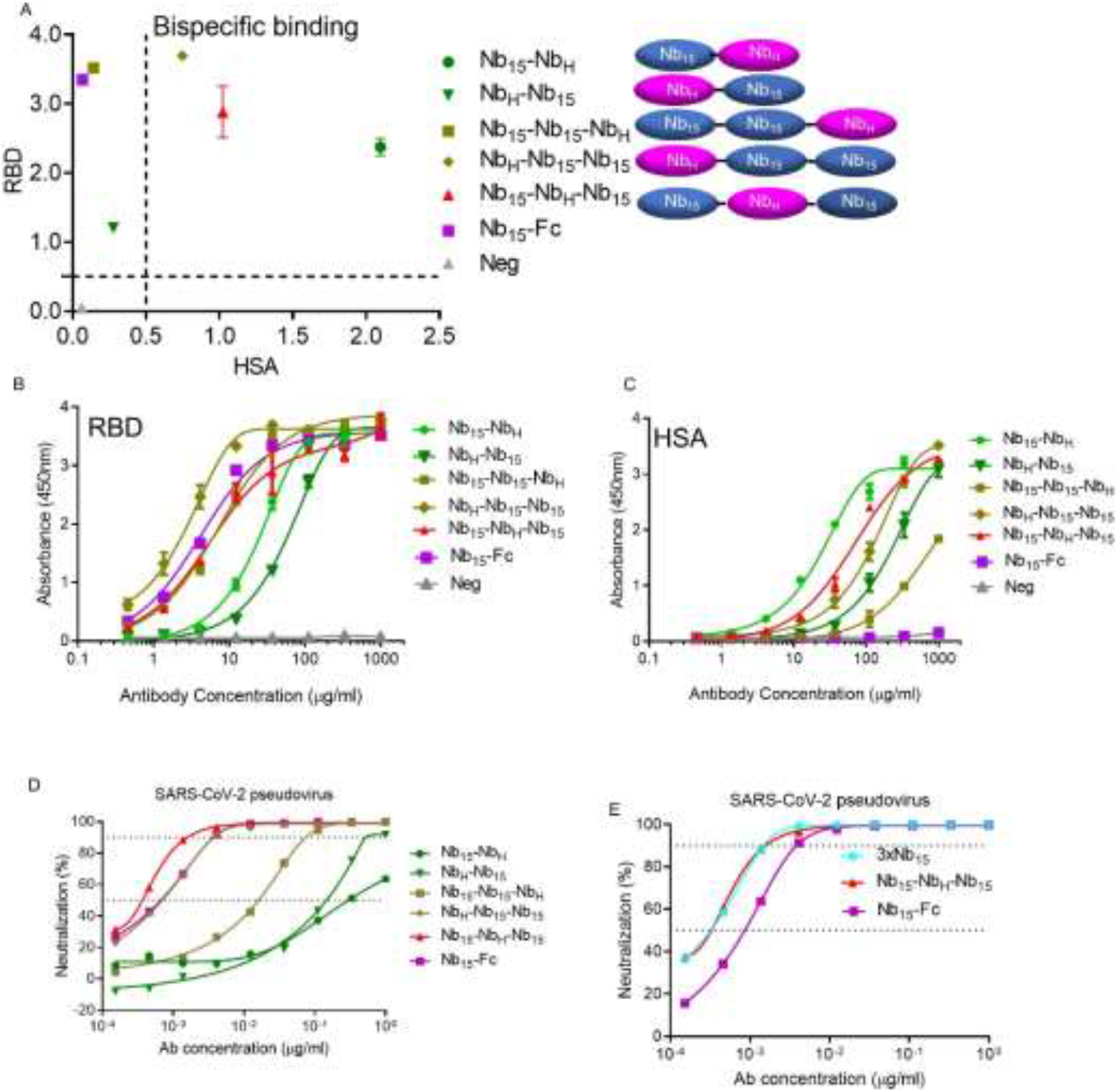
Design and characterization of bispecific Nbs. (**A**) Various Nbs at 37 μg/ml binding to RBD and HSA protein identified by ELISA. The binding curve of Nbs interacting with RBD protein (**B**) and HSA protein (**C**) identified by ELISA. (**D**) The neutralizing potency of bispecific Nbs against SARS-CoV-2 pseudovirus measured by neutralization assay. (**D**) The neutralizing potency of Nb_15_-Fc, 3×Nb_15_ and Nb_15_-Nb_H_-Nb_15_ against SARS-CoV-2 pseudovirus infection measured by neutralization assay. Data represent as mean ± SEM. All experiments were repeated at least twice.

### Pharmacokinetics and delivery of Nbs constructs

Given 3×Nb_15_, Nb_15_-Nb_H_-Nb_15_ and Nb_15_-Fc exhibited comparable neutralization activity (Fig. 2E), these three constructs were evaluated for their *in vivo* pharmacokinetic activity, and the results showed that, when administrated via *i.n., i.p.*, or *i.v*, 3×Nb_15_ was rapidly metabolized as compared to Nb_15_-Fc and Nb_15_-Nb_H_-Nb_15_ (Fig. 3A-C); therefore, 3×Nb_15_ was ruled out for further analysis. To determine the tissue distribution of Nbs, YF®750 SE-labeled Nb_15_-Nb_H_-Nb_15_ (Nb_15_-Nb_H_-Nb_15_-YF750) were administered via *i.n.*, *i.p*. or *i.v.* in mouse model. The results revealed that the fluorescence in trachea could be detected only when Nb_15_-Nb_H_-Nb_15_-YF750 administered *i.n*.. Furthermore, The fluorescence intensity was higher in lungs when Nb_15_-Nb_H_-Nb_15_-YF750 was administered *i.n*. (6.9 × 10^10^ ph/s) than that when administered *i.p*. or *i.v.* (1.4 × 10^10^ and 4.3 × 10^10^ ph/s, respectively) (Fig. 3D and 3E). In addition, the results also showed that Nb_15_-Nb_H_-Nb_15_ could reach lungs, and sustained for more than 168 h (7 d) when administrated *i.n*.; in contrast the fluorescence could only be detected between 1 and 2 h post *i.p*. infusion (Fig. 3F and 3G). These results suggest that *i.n.* administration of Nb_15_-Nb_H_-Nb_15_ will be a favorable route for antibody to reach nasopharynx and lungs where SARS-CoV-2 replicates. Therefore, to avoid the potential ADE associated by Fc in the Nb-Fc, we selected Nb_15_-Nb_H_-Nb_15_ for further efficacy evaluation *in vivo*.

**Figure 3.**
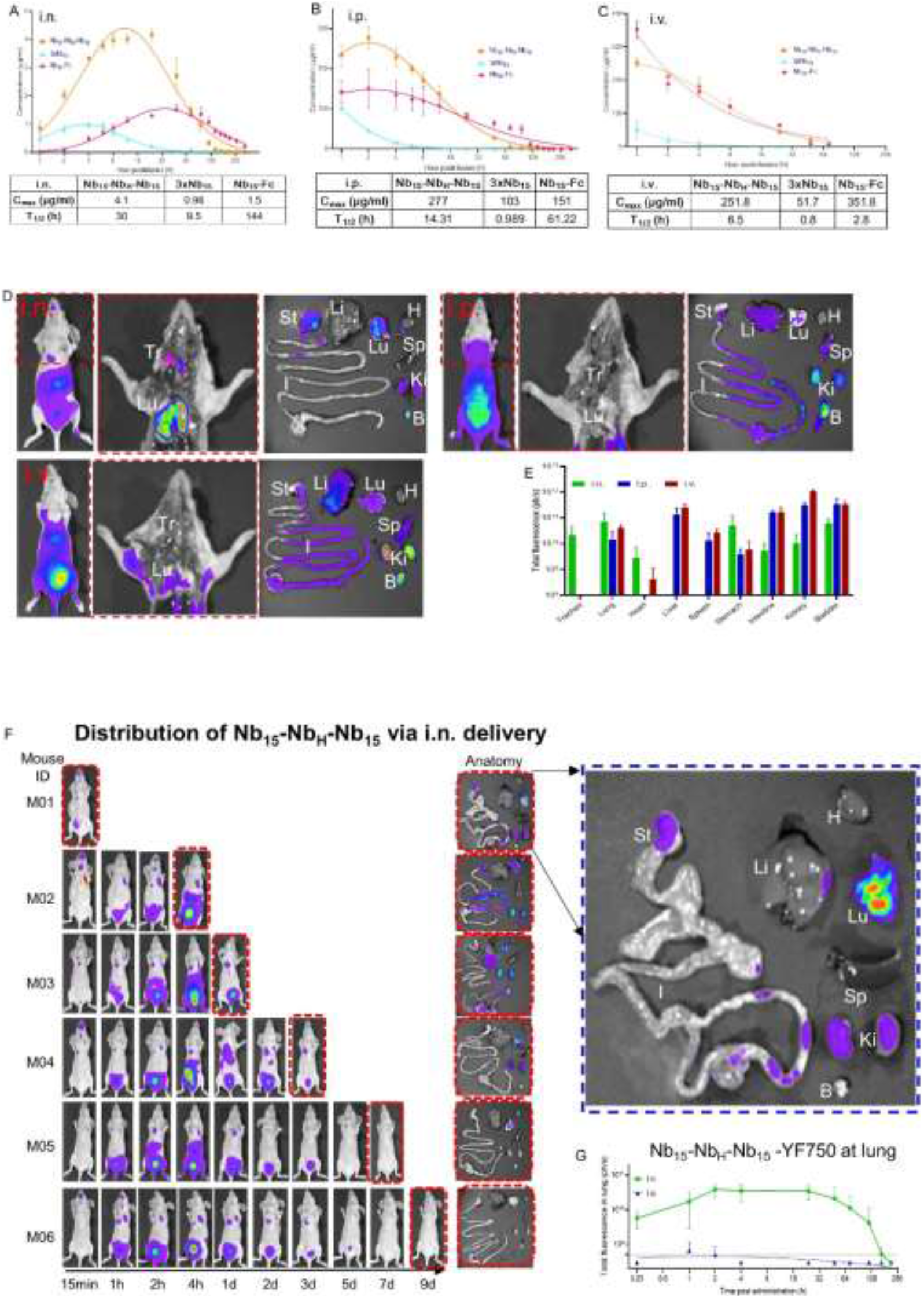
Pharmacokinetics of Nb_15s_ *in vivo*. Bioavailability and T_1/2_ (half-life) of Nb_15s_ in BALB/c mice. Nb_15_ variants were intranasally (i.n.) administered into mice (n=3, Female) at 200 ug (average of 10 mg/kg mice) (**A**), intraperitoneally (i.p.) administered into mice (n=3, Female) at 400 ug (average of 20 mg/kg mice) (**B**), intravascularly (i.v.) administered into mice (n=3, Female) at 400 ug (average of 20 mg/kg mice) (**C**), respectively. Serum concentrations of the Nbs were determined at indicated time points by ELISA. Nb_15_ variants are colored as follows; Nb_15_-Fc (red), Nb_15_-Nb_H_-Nb_15_ (orange) and 3×Nb_15_ (cyan). C_max_, maximum observed plasma concentration, T_1/2_, time of half-life. Data represent as mean ± SEM. (**D**) Spatial distribution of Nb_15_-Nb_H_-Nb_15_YF750 1 hour after infusion into mice (n=3 in each group) via i.n., i.p. and i.v. was detected by NightOwl LB 983. The middle figure in red dash line is the dissected image of the left mouse in the red dash line. The right figure is the Organs from dissected mice which were imaged immediately after sacrifice. Tr, Trachea; Lu, Lung; H, Heart; Li, Liver; Sp, spleen; St, Stomach; I, Large and small intestine; Ki, Kidneys; B, Bladder.(**E**) The fluorescence intensity (ph/s) summary of each organ in **D** was quantified and presented as the mean ± SEM. (**F**) Pharmacokinetic of Nb_15_-Nb_H_-Nb_15_-YF150 via intranasal administration at indicated time point. Mice were sacrificed at the indicated time point for the analysis of fluorescence intensity in various organs labeled as D. The blue dash line figure is the enlarged image of the individual figure indicated by corresponding arrows. (**G**) Nude mice (n=3-6) were administered with Nb_15_-Nb_H_-Nb_15_ – YF750 i.n. or i.p.. The fluorescence intensity at the lung location as the yellow dash line circle of M02 in **F** was measured at the indicated time point. Data represent as mean ± SEM.

### *In vitro* characterization of Nb_15_-Nb_H_-Nb_15_

Nb_15_-Nb_H_-Nb_15_ was further characterized *in vitro*. Nb_15_-Nb_H_-Nb_15_ exhibited specific binding to RBD and HSA with *K*_*D*_ values of 0.54 and 7.7nM, respectively. In addition, Nb_15_-Nb_H_-Nb_15_ also showed specific binding with murine serum albumin (MSA) with *K*_*D*_ values of 14.5 nM, indicating that mice can be used as an animal model to investigate the half-life of Nb_15_-Nb_H_-Nb_15_ (Fig. 4A-B and Fig. S8). Furthermore, Nb_15_-Nb_H_-Nb_15_ exhibited sub-ng/ml (pM) potency against both the wild-type and currently circulating mutant variants of SARS-CoV-2 (Fig. 4C and Table S2). Importantly, Nb_15_-Nb_H_-Nb_15_ showed comparable potency against the SARS-CoV-2 variants with D614G and N501Y mutations that circulate predominantly in the UK and South Africa. D614G and N501Y variants conferred enhanced replication and transmissibility and emerged as the predominant global variants with high transmission^30^. Nb_15_-Nb_H_-Nb_15_ also showed excellent thermal stability by retaining 100% and 83% activities even at 70 °C and 80 °C for one hour, respectively (Fig. 4D-E and Table S4). Furthermore, Nb_15_-Nb_H_-Nb_15_ retained 100% activity after aerosolization, indicating the potential application as a nebulized drug (Fig. 4D-E and Table S4).

**Figure 4.**
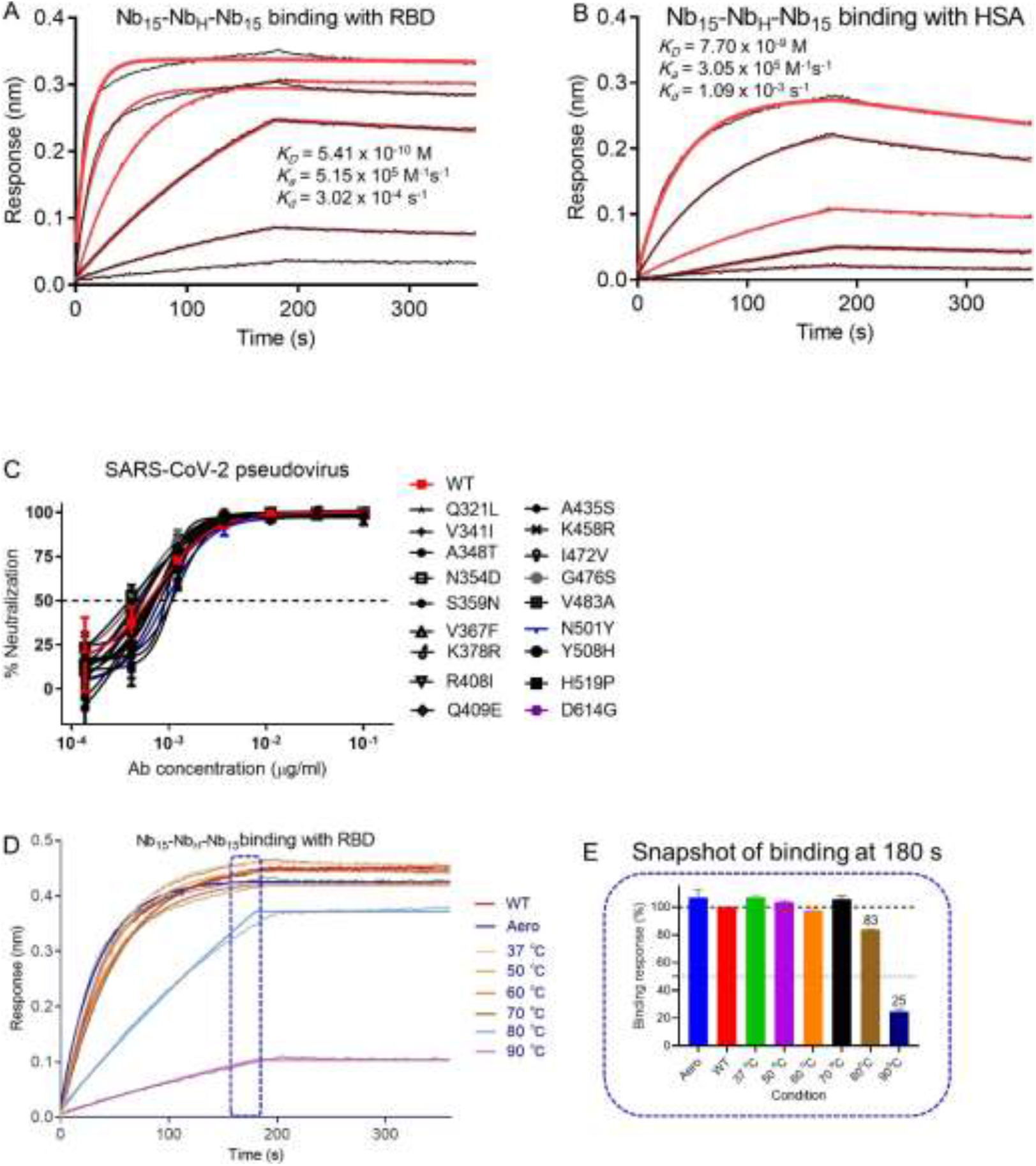
Functional characterization of Nb_15_-Nb_H_-Nb_15_. Kinetic binding curve of Nb_15_-Nb_H_-Nb_15_ at the concentration 300 nM, 100nM, 33.3 nM,11.1nM, 3.7nM and 1.2 nM with RBD (**A**) and HSA (**B**), respectively, by BLI. Binding curves are colored black, and fit of the data to a 1:1 binding model is colored red. (**C**) The neutralization curve of Nb_15_-Nb_H_-Nb_15_ inhibiting SARS-CoV-2 pseudovirus and its variants with amino acid point mutation as indicated. Data represent as mean ± SEM. (**D**) Binding curve of RBD with Nb_15_-Nb_H_-Nb_15_ at the concentration of 133 nM (5μg/ml) before ( no treatment, WT) or after aerosolization (Aero) or after treatment at the indicated temperature, including 37 °C, 50 °C, 60 °C, 70 °C, 80 °C and 90 °C for one hour. Binding curves are colored black, and fit of the data to a 1:1 binding model is colored as indicated. (**E**) Snapshot of the relative binding response of the highest binding response at the indicated condition /the highest response of RBD with Nb_15_-Nb_H_-Nb_15_ in no treatment condition (WT). The relative binding of WT was normalized as 100%. Data represent as mean ± SEM. All experiments were repeated at least twice.

### *In vivo* anti-SARS-CoV-2 activity of Nb_15_-Nb_H_-Nb_15_

To evaluate the efficacy of Nb_15_-Nb_H_-Nb_15_ *in vivo*, hACE2 transgenic mice were challenged with SARS-CoV-2, and Nb_15_-Nb_H_-Nb_15_ was administrated *i.n.* either before or after viral challenge for prophylactic or therapeutic efficacy (Fig. 5A). Viral RNA was detected in lungs in the control mice (6.28 × 10^5^ copies/mg on average in SARS-CoV-2 group, n=5) and the isotype treated control mice (7.8 × 10^4^ copies/mg on average in isotype group, n=3,). For the prophylactic group, no viral RNA or infected cells was detected in 100% (5/5) of the mice when 250 μg (average of 10 mg/kg) Nb_15_-Nb_H_-Nb_15_ was administrated via *i.n*. 24 hours before SARS-CoV-2 infection (Nb_15_-Nb_H_-Nb_15_ −24h group, n=5), as evidenced by real-time PCR and immunofluorence staining (Fig. 5B-D). 100% mice were also completely protected when 250 μg Nb_15_-Nb_H_-Nb_15_ were administrated via *i.n.* 1 hour postinfection, as no viral RNA and infected cells were detected in all infected mice (Nb_15_-Nb_H_-Nb_15_ 1h group, n=5) (Fig. 5B-D). Significantly lower SARS-CoV-2 RNA copies (9.98 ×10^3^ copies/mg on average) were detected in the lungs of the mice treated with Nb_15_-Nb_H_-Nb_15_ *i.n*. 24 h postinfection (Nb_15_-Nb_H_-Nb_15_ 24 h group, n=5) than that in the control mice (6.28 × 10^5^ copies/mg on average in SARS-CoV-2 group and 7.8 × 10^4^ copies/mg in isotype control) (Fig. 5B-D). Nb_15_-Fc inhibited viral replication and reduced the viral copies number (average of 7.59 ×10^3^ copies/mg in Nb_15_-Fc 1h group, n=5) but failed to provide complete protection under the same condition as Nb_15_-Nb_H_-Nb_15_ (Fig. 5B-D). Furthermore, histopathological analysis of lung tissues showed that SARS-CoV-2 challenge induced severe lung lesions, as shown by the infiltration of inflammatory cells and thickened alveolar septa (Fig. 5D). In contrast, the lungs of the mice receiving Nb_15_-Nb_H_-Nb_15_ or Nb_15_-Fc treatment showed no apparent pathological changes (Fig. 5D). Together, Nb_15_-Nb_H_-Nb_15_ at an average of 10 mg/kg via *i.n*. administrated 24 h before or 1 h after challenge provided complete protection against SARS-CoV-2 infection, and significantly inhibited SARS-CoV-2 replication when the antibody was administrated 24 h postinfection. Nb_15_-Fc used at an average of 10 mg/kg via *i.n*. administrated 1 h after challenge significantly reduced viral load but failed to provide complete protection. We noted that those mice receiving Nb_15_s treatment showed less weight loss than the control mice but did not achieve statistical difference (Fig. 5E-F). These results indicate that Nb_15_-Nb_H_-Nb_15_, when used early during infection, confered higher protection efficacy than used at later time point. In summary, the Nb_15_-Nb_H_-Nb_15_ configuration administered via *i.n*. was superior to Nb_15_-Fc and exhibited both prophylactic and therapeutic efficacy against SARS-CoV-2 challenge.

**Figure 5.**
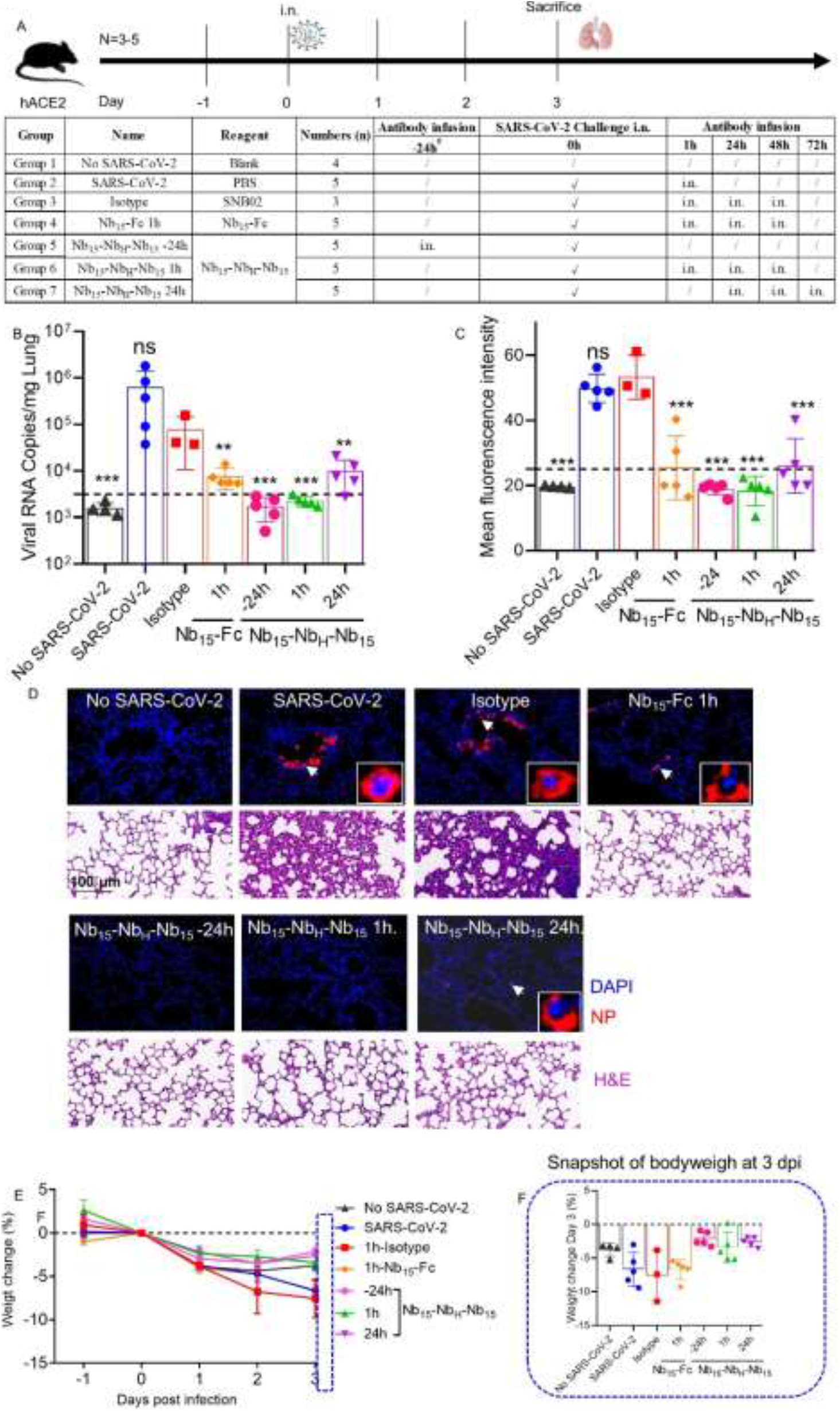
The efficacy of Nb_15s_ evaluated in hACE2 transgenic mice challenged by SARS-CoV-2. (**A**) Experimental schedule of Nb_15s_ in the prevention and treatment of SARS-CoV-2 infection. The below table summary of groups (n=3-5 mice) with different treatment. (**B**) Viral loads in lung among 7 groups were measured by qRT-PCR. The name of each group in *X* axis was indicated as the table in **A**. Each dot represents one mouse. The limit of detection was 3160 copies/mg referenced to blank control (No SARS-CoV-2 group). (**C**) Sections of lung were analyzed by immunofluorescence staining using antibodies specific to SARS-CoV-2 NP in red and DAPI for nuclei in blue, respectively. The fluorescence signal intensity of red was taken as a quantitative indicator for viral infection, which was calculated by ImageJ software. (**D**) Representative sections of lung in **C** were visualized under the × 20 objective. The insets are enlarged images of individual cells indicated by corresponding arrows. H&E staining was conducted to analyze the lung inflammation and observed at the indicated scale bar. (**E**) Body weight of mice among the above 7 groups were recorded. Each line represents data from one group. (**F**) Snapshot of body weight on 3 days post infection in (**E**) was plotted. Data represent mean ± S EM; One-way or two-way ANOVA were performed to compare treatment group with the isotype control group. ns, no significance; **, *P* < 0.01, ***, *P* < 0.001. Data of B, C, E and F represent as mean ± SEM. All experiments of B and C were repeated twice.

## Discussion

In this study, three potent neutralizing Nb-Fcs were isolated from a phage display platform derived from an SARS-CoV-2 S protein immunized alpaca. These three RBD-specific Nb-Fcs exhibited potent inhibitory activities against 15 mutant variants of SARS-CoV-2 at ng/ml concentration (Fig. 1E-G and Table S2). The IC_50_ values are comparable to those of the most potent neutralizing antibodies, or Nbs reported^4,21,22,27,28^. These 15 representative variants of SARS-CoV-2 are identified to represent over 7000 unique viral genomes ^2^. More importantly, recently arising variants with D614G mutation were also sensitive to the neutralization by Nb_15_-Fc, Nb_22_-Fc and Nb_31_-Fc with similar sensitivity (Fig. 1E-G and Table S2). These evidences demonstrate the broadly neutralizing activity of these Nbs against SARS-CoV-2 and suggest that the Nb-Fcs target at a highly conserved epitope on RBD protein.

To improve potency, prolong *in vivo* half-life and avoid potential Fc-mediated ADE, Nb_H_ specific for HSA and MSA was used to construct a trimeric Nb_15_ (Nb_15_-Nb_H_-Nb_15_) for SARS-CoV-2, and the resulting Nb_15_-Nb_H_-Nb_15_ exhibited the highest neutralization potency with the IC_50_ value of 0.4 ng/ml among other configurations, including Nb_15_-Nb_H_, a configuration reported earlier^29^. Interestingly, we found that Nb_15_-Nb_H_ and Nb_H_-Nb_15_ with the same components exhibited distinct neutralization potencies with IC_50_ values of 552.3 ng/ml and 197.4 ng/ml. In addition, Nb_H_-Nb_15_-Nb_15_ and Nb_15_-Nb_15_-Nb_H_ also displayed distinct neutralization potencies with IC_50_ values of 25.1 ng/ml and 8 ng/ml (Fig. 2D and Table S3), indicating that Nb configuration has impact on the neutralizing activity. In addition, heterotrimeric bispecific configuration is superior to the bispecific heterodimer. We also noted that the bi-, tri- and tetra-valent configurations exhibited comparable potency with IC_50_ values of 2.8, 3.5 and 2.3 ng/ml (11, 9.0, 4.3 pM), respectively. The neutralizing potency did not correspond to the valence increase when there are two or more than two Nb_15_s though monomeric 1×Nb_15_ had much lower inhibitory activity (Fig. S7C and Table S3). Noted that Nb_15_-Nb_H_-Nb_15_ shows higher potency than Nb_H_-Nb_15_-Nb_15_, Nb_15_-Nb_15_-Nb_H_, and all the homomultimers, suggesting that the position of Nb_H_ plays important roles in neutralization activity. We speculate that in Nb_15_-Nb_H_-Nb_15_ Nb_H_ may space out the two Nb_15_s to either avoid cross intereference with each other or allow better binding of the trimeric Nb to S proteins on the viral particle. Furthermore, Nb_15_-Nb_H_-Nb_15_ displayed comparable neutralizing potency as those of Nb_15_-Fc and 3×Nb_15_, and higher neutralization potency and longer half-life than 3×Nb_15_ *in vivo* when delivered via *i.n*. , *i.p*. or *i.v*.. Importantly, Nb_15_-Nb_H_-Nb_15_ exhibited broadly neutralizing activities against all the SARS-CoV-2 variants that we tested, including those with D614G and N501Y mutations that currently circulate in the UK and South Africa. These mutant viruses conferred enhanced replication, transmissibility and emerged as the global predominant circulating variants and are great public health concerns^30^.

Though several neutralizing antibodies against SARS-CoV-2 are in various stages of the clinical trials or have been approved as an emergency therapy; most of these antibodies are of limited efficacy. To the best of our knowledge, our study is the first to demonstrate the efficacy of Nb for both the prevention and treatment of SARS-CoV-2 infection (Fig. 5). SARS-CoV-2 is mainly present in the nasopharynx and lungs^31,32^. Differing from many previously reported systemic delivery route, Nb_15_-Nb_H_-Nb_15_ was delivered to the site of infection. Direct administration to the airways is likely to provide faster and more robust antiviral activity in the respiratory tract, where the virus gains entry and replicate ^16,32^, as intranasal delivery has been shown to result in fast and efficient drug delivery to the main site of SARS-CoV-2 infection, i.e., the upper and lower respiratory tract^32^. Indeed, the therapeutic effect of topical administration of ALX-0171 displayed promising results in reducing the RSV viral load^14,16^. To our best knowledge, the current study is the first to evaluate the *in vivo* efficacy of Nbs against SARS-CoV-2 infection via intranasal delivery.

In summary, comparing to the configurations of Nb-Nb_H_, Nb-Fc and Nb homotrimer, a novel construct of Nb_15_-Nb_H_-Nb_15_ exhibited higher neutralization activity and longer half life *in vivo*. Nb_15_-Nb_H_-Nb_15_ exhibited highly potent antiviral activity with broad specificity against a large panel of SARS-CoV-2 clinical variants. Furthermore, direct delivery of Nb_15_-Nb_H_-Nb_15_ to the airways/lungs by intranasal route proved an effective mode of drug delivery, and the outstanding thermal stability of Nb_15_-Nb_H_-Nb_15_ is an additional advantage. We suggest that respiratory delivery of Nb_15_-Nb_H_-Nb_15_ is a promising route for the prevention and treatment of SARS-CoV-2 infection, and thus warrants further clinical evaluation.

## Materials and Methods

### 1. Alpaca immunization

250 μg the extracellular domain of SARS-CoV-2 spike protein fused with His tag (S1+S2 ECD, S, cat.# 40589-V08B1, Sino Biological) was emulsified with 250 μl Freund’s complete adjuvant (F5881-10ML, Sigma) to immunize an alpaca. On day 14 and 28, the alpaca was boosted twice with 250 μg S protein in 250 μl Freund’s incomplete adjuvant (F5506-10ML, Sigma). One week following the 2^nd^ immunization, we collected the blood samples to measure anti-serum titer. One week after the 3^rd^ immunization, 100 ml of blood was collected to measure anti-serum titer and construct a phage library displaying Nb.

### 2. SDS-PAGE and Western blotting (WB)

The purified protein or antibody was separated by electrophoresis in a 7.5%-12% polyacrylamide gel. The separated protein or antibody was revealed either using Coomassie blue or transferred to PVDF membrane for WB analysis under reducing or non-reducing conditions with β-mercaptoethanol. The membrane was first blocked and then incubated overnight at 4 °C or 37 °C for one hour with diluted plasma or antibody, followed by incubation with the secondary antibody of either anti-human IgG or anti-rabbit IgG conjugated with an IRDye 800CW (cat.# 926-32232, Rockland). Protein bands were visualized using the Odyssey Image System (Li-COR).

### 3. ELISA analysis

Anti-sera titer and antibody characterization or antibody quantification *in vivo* were examined by ELISA as reported in our previously published method^33^ with modifications. In brief, the protein was coated to high protein-binding ELISA plates (Corning) at a concentration of 0.5 μg/ml, 100 μl per well at 37 °C for 2 hours (h) or 4 °C overnight. After washing, blocking buffer with 5% non-fat milk in PBS was added and incubated at 37 °C for 1 h. After washing 2-4 times, 100 μl serially diluted anti-serum or purified antibody was added and incubated at 37 °C for 1.5 h. Following washing, goat anti-llama IgG (H+L) secondary antibody with HRP (Novus, cat.# NB7242, 1:10000 dilution) was added and incubated at 37 °C for 1 h. Accordingly, 3,3′,5,5′-Tetramethylbenzidine (TMB, Sigma) substrate was added at 37 °C for 10 minutes (min); and the reaction was stopped by adding 10 μl 0.2 M H_2_SO_4_. The optical densities at 450 nm were measured using the Infinite 200 (Tecan, Ramsey, MN, USA). Antibody titers were defined as the highest dilution when the diluted serum produced at least 2.1-fold optical density readout as compared to the control serum sample at the same dilution.

### 4. Construction of a phage library displaying Nbs

Nb phage library was constructed following our previously published method with some modifications^13^. In brief, PBMCs were isolated from 100 ml blood of immunized alpaca using a lymphocyte separation solution (cat.# 17-1140-02, Ficoll-Paque Plus, GE). RNA was extracted and reverse transcribed into cDNA by oligo (dT) and random hexamers as primers using the TRIzol kit (cat.# 15596018, Ambio By Life Technologies), following manufacturer’s instruction. The alpaca Nb gene was amplified with the combination of primers and cloned into phV1 phagemid plasmid (Y-Clone, Ltd., China) to transform TG1 bacteria.

### 5. Panning Nb phage library and phage ELISA

Affinity selection for S-binding recombinant phages was performed as previously reported with the following modifications^34^. The Nb-phagemid-transformed bacteria were rescued with M13KO7 helper phage (cat.# 18311019, Invitrogen), and precipitated with PEG/NaCl. The phage Nb antibody library was enriched three times with 50 μg/ml of S protein. The enriched phage was eluted, transformed, and selected for the monoclonal phage to be evaluated by phage ELISA.

### 6. Phage ELISA

200 ng S or RBD protein in coating buffer (pH 9.6) was used to coat 96-well plates (cat.# 9018, Corning) at 4 °C overnight. After washing, the plates were blocked with blocking buffer (3% BSA in PBST) for 1 h at 37 °C, and then incubated with library phages or single clone phage in bacterial supernatant at 4 °C for 1.5 h. After washing, an anti-M13 bacteriophage antibody with HRP (1:10000 dilution, cat.# 11973-MM05T-H, Sino Biological) was added and incubated at 37 °C for 1 h. Accordingly, TMB substrate(Sigma) was added at 37 °C for 10 min; 10 μl 0.2 M H_2_SO_4_ was added to stop the reaction. Optical densities were measured at 450 nm using the Infinite 200 (Tecan, Ramsey, MN, USA). Clones with readout at 450 nm >0.5 were sequenced.

### 7. Expression and purification of Nbs with different formats

To facilitate the purification and prolong the half-life of the Nb antibody, the Fc1 gene (CH2-CH3) of the human monoclonal antibody was fused with the Nb gene (Nb-Fc), as our previously published method^13^. In addition, to improve the activity of Nb, we constructed Nbs with various configurations wherein (GGGGS)_3_ linkers were introduced between Nbs in dimeric and trimeric forms. To facilitate protein purification, a 6xHis-tag was fused to the N terminus of the Nbs of monomeric, dimeric or trimeric configuration. The Nbs with different configurations were finally cloned into the pCDNA3.4 eukaryotic expression vector (Invitrogen), which were transfected into 293F cells to produce Nbs with different configurations. Nb fused with Fc, or His tag was purified using Protein G (cat.# 20399, Thermo Scientific) and Ni-NTA (cat.# R901100, Thermo Fisher Scientific), respectively.

### 8. Neutralization activity of Nbs against pseudovirus

Pseudovirus neutralization assay was performed as previously described with the following modifications^4^. SARS-CoV-2, SARS-CoV, and MERS-CoV pseudoviruses were produced by co-transfection of pNL4-3.Luc.R-E-, an HIV-1 NL4-3 luciferase reporter vector that contains defective Nef, Env and Vpr (HIV AIDS Reagent Program), and pCDNA3.1 (Invitrogen) expression vectors encoding the respective spike proteins (MN988668.1 for SARS-CoV-2, AAP13567.1 for SARS-CoV, AFS88936.1 for MERS-CoV) into 293T cells (ATCC). Pseudovirus containing supernatants were collected after 48 h, and viral titers were measured by luciferase assay in relative light units (Bright-Glo Luciferase Assay Vector System, Promega Biosciences). S genes of SARS-CoV-2 variants with indicated mutations based on the human codon optimized S gene (Accession number: MN988668.1) were synthesized, and the corresponding pseudoviruses were produced following above protocol. For neutralization assay, SNB02 (Nb-Fc) against SFTSV^13^ served as a control. Neutralization assays were performed by incubating pseudoviruses with serial dilutions of purified Nbs or serum at 37 °C for 1 h. HEK293T-ACE2 cells (cat.# 41107ES03, Yeasen Biotech Co., Ltd. China) for SARS-CoV-2 and SARS-CoV, Huh7 cells (ATCC) for MERS-CoV (approximately 1.5×10 ^4^ per well) were then added in duplicate to the virus-antibody mixture. Half-maximal inhibitory dilution (ND_50_) of the evaluated sera or half-maximal inhibitory concentrations (IC_50_) of the evaluated Nbs were determined by luciferase activity 48 h after exposure to virus-antibody mixture, and analyzed by GraphPad Prism 8.01 (GraphPad Software Inc.).

### 9. Neutralization activity of Nbs against live SARS-CoV-2

SARS-CoV-2 focus reduction neutralization test was performed in a certified Biosafety Level 3 laboratory, as previously described with the following modifications^6^. Briefly, a clinical isolate (Beta/Shenzhen/SZTH-003/2020, EPI_ISL_406594 at GISAID) previously obtained from a nasopharyngeal swab of an infected patient was used for the analysis. Serial concentrations of Nbs were mixed with 75 μl of SARS-CoV-2 (8×10 ^3^ focus forming unit/ml, FFU/ml) in 96-well microwell plates and incubated at 37 °C for 1 h. The mixtures were then transferred to 96-well plates seeded with Vero E6 cells and incubated at 37 °C for 1 h. Next, the inoculums were removed prior to the addition of the overlay media (100 μl MEM containing 1.6% carboxymethylcellulose, CMC) and the plates were then incubated at 37 °C for 24 h. Cells were fixed with 4% paraformaldehyde solution for 30 min, and then the overlays were removed. Cells were permeabilized with 0.2% Triton X-100 and incubated with cross-reactive rabbit anti-SARS-CoV-N IgG (Sino Biological, Inc) for 1 h at room temperature before the addition of HRP-conjugated goat anti-rabbit IgG (H+L) antibody (Jackson ImmunoResearch) and further incubated at room temperature. The foci were stained with KPL TrueBlue Peroxidase substrates (SeraCare Life Sciences Inc.) and were counted with an EliSpot reader (Cellular Technology Ltd.).

### 10. Affinity determination by Bio-Layer Interferometry (BLI)

Affinity assays were performed on a ForteBio OctetRED 96 biolayer interferometry instrument (Molecular Devices ForteBio LLC, Fremont, CA) at 25 °C with shaking at 1,000 rpm. To measure the affinity of Nbs with human Fc tag, anti-human Fc (AHC) biosensors (cat.# 18-5060, Fortebio) were hydrated in water for 30 min prior to 60 seconds (sec) incubation in a kinetic buffer (PBS, 0.02% (v/v) Tween-20, pH 7.0). Either Nb-Fc in cell supernatant or purified Nb-Fcs were loaded in a kinetic buffer for 200 sec prior to baseline equilibration for 200 sec in a kinetic buffer. Association of SARS-CoV-2 RBD in a two-fold dilution series from 20 nM to 2.5 nM was performed prior to dissociation for 180 sec. To measure the affinity of Nbs without Fc tag, RBD protein was coupled to AR2G biosensor (cat.# 18-5092, Fortebio) via BLI instrument according to the instructions of the amino coupling kit. Association of Nbs in a serial dilution was performed prior to dissociation for 180 sec. After each cycle, the biosensors were regenerated via 3 short pulses of 5 sec each of 100 mM pH 2.7 glycine-HCL followed by running buffer. The data were baseline subtracted before fitting performed using a 1:1 binding model and the ForteBio data analysis software. *K*_*D*_, *Ka* and *Kd* values were evaluated with a global fit applied to all data.

### 11. Epitope binning by BLI

The epitope binning assay was performed with AR2G biosensor (cat.# 18-5092, Fortebio) following the manufacturer’s protocol ‘in-tandem assay’ as previously reported^4^. After loading the RBD protein, a saturating concentration of antibody or Nbs (50 μg/ml) as the first antibody was added for 300 sec following with the baseline step with 30 s immersion in 0.02% PBST. The second competing concentration of antibody or Nb (50 μg/ml) was then added for 300 sec to measure binding in the presence of the first saturating antibody or Nb. GraphPad was used to illustrate the time-response course of two antibodies binding to RBD protein.

### 12. Evaluating the efficacy of Nbs in SARS-CoV-2 infected hACE2 mice

A total of 31 8-week-old male transgenic hACE2 mice (C57BL/6J) (cat.# T037630, GemPharmatech Co., Ltd., Nanjing, China) were challenged with SARS-CoV-2 as previously reported^35^ with following modifications. The mice were split into seven groups (n=3-5) for either prophylactic or therapeutic evaluation, as described in Fig. 5A. Mice without any challenge and treatment served as blank control (No SARS-CoV-2, n=4). Mice challenged with SARS-CoV-2 were taken as infection control (SARS-CoV-2, n=5). 250 μg SNB02 (Y-Clone, China), an anti-SFTSV antibody constructed by Nb fused with human Fc1 (Nb-Fc)^13^, was intranasally injected 1 h after infection and served as an isotype treated control (Isotype). For the prophylactic group, mice were intranasally injected with Nb_15_-Nb_H_-Nb_15_ at a dose of 250 μg/mouse (average of 10 mg/kg) 24 h before infection (Nb_15_-Nb_H_-Nb_15_ −24h, n=5). For the therapeutic group, mice were intranasally injected with N Nb_15_-Nb_H_-Nb_15_ at a dose of 250 μg/mouse (average of 10 mg/kg) 1 h or 24 h after infection (named as Nb_15_-Nb_H_-Nb_15_ 1h and Nb_15_-Nb_H_-Nb_15_ 24 h, n=5, respectively). As a comparison, Nb_15_-Fc at a dose of 250 μg/mouse (average of 10 mg/kg) was intranasally injected 1 h after infection (Nb_15_-Fc 1 h). Body weight of every mouse was measured daily. Transgenic hACE2 mice typically clear virus within five days after SARS-CoV-2^35^. Accordingly, the mice were sacrificed at 3 days post infection (dpi), and the lungs were collected for viral load determination and tissue sections for hematoxylin and eosin (H&E) and immunofluorescence staining.

### 13. Viral load measurement by quantitative RT-PCR

Viral load was detected by quantitative real-time PCR (qRT-PCR) on RNA extracted from the supernatant of lung homogenates as described previously^36^. Briefly, lung homogenates were prepared by homogenizing perfused whole lung using an electric homogenizer. The supernatant was collected, and total RNA was extracted. Each RNA sample was reverse transcribed to 50 μl cDNA with RT-PCR Prime Script Kit (Takara). The cDNA (5 μl) was used in a 25 μl qRT-PCR reaction with the TaqMan Universal PCR Master Mix (Life Technologies), a TaqMan probe (5′-FAM− CAGGTGGAACCTCATCAGGAGATGC −MGB-3′), and primers designed to target the orf1ab gene of SARS-CoV-2 (5′-GTGARATGGTCATGTGTGGCGG -3′ and 5′-CARATGTTAAASACACTATTAGCATA -3′). The samples were run in triplicate on an ABI 7900 Real-Time System (Applied Biosystems, Thermo Fisher Scientific). The following cycling conditions were used: 1 cycle of 50 °C for 2 min, 1 cycle of 95 °C for 10 min, and 40 cycles of 95 °C for 15 sec and 58 °C for 1 min. The virus titer was determined by comparison with a standard curve generated using RNA extracted from a serially diluted reference viral stock. All experiments were performed in a Biosafety Level 3 facility.

### 14. Immunofluorescence staining of SARS-CoV-2-infected cells in tissues

Lung tissues were immersed in 10% neutral buffered formalin (cat.# Z2902, Sigma) for 24 h. After the formalin fixation, the tissues were placed in 70% ethanol (Merck) and subsequently embedded with paraffin. Tissue sections (4-μm thick) were used for immunofluorescence staining for SARS-CoV-2 detection using the Coronavirus nucleocapsid antibody (cat. 40143-MM05, Sino Biological). Images were obtained by OLYMPUS IX73 using HCImage Live (×64) software and analyzed by ImageJ (NIH).

### 15. Pharmacokinetics of Nbs *in vivo*

Purified Nbs were injected intranasally (*i.n*.), intraperitoneally (*i.p*.) or intravascularly into BALB/c (Qing Long Shan Animal Center, Nanjing, China) at a dose of 10-20 mg/kg. ELISA was used to measure the serum concentration of Nbs. The T_1/2_ of Nbs was computed as ln (2)/k, where k is a rate constant expressed reciprocally of the × axis time units by the one phase decay equation or plateau followed one phase decay in the GraphPad software.

### 16. Spatial distribution of Nbs *in vivo*

Nbs were labeled with far infrared dye YF®750 SE (US EVERBRIGHT INC, YS0056) (named as Nbs-YF750). Purified Nbs-YF750 were injected *i.n., i.p*. or *i.v.* into nude mice (18-22g, Qing Long Shan Animal Center, Nanjing, China) at a dose of 10-20 mg/kg. Images were observed at Ex:740 nm/Em:780 nm by NightOWL LB 983 (Berthold, Germany) at the indicated time point. Images were analyzed using Indigo imaging software Ver. A 01.19.01.

### 17. Statistics

All statistical analyses were performed using GraphPad Prism 8.01 software (GraphPad) or OriginPro 8.5 software (OriginLab). ANOVA was performed for group comparisons. P < 0.05 was considered as statistically significant with mean ±SEM or mean ±SD.

### 18. Study approval

The study and the protocol for this research were approved by the Center for Public Health Research, Medical School, Nanjing University. All animal experimental procedures without infection were approved by the Committee on the Use of Live Animals by the Ethics Committee of Nanjing University. All of the animals infected by SARS-CoV-2 were handled in Biosafety Level 3 animal facilities in accordance with the recommendations for care and use of the Institutional Review Board of Wuhan Institute of Virology of the Chinese Academy of Sciences (Ethics Number: WIVA11202003). All the authors declare their compliance with publishing ethics.

## Acknowledgments

This work was supported by National Science Foundation of China (NSFC) (No. 81803414, 31970149), the Major Research and Development Project (2018ZX10301406), Nanjing University-Ningxia University Collaborative Project (Grant# 2017BN04), Jiangsu Province Natural Science Foundation for Young Scholar (Grant# BK20170653), Key Natural Science Foundation of Jiangsu Province (Grant# ZDA2020014), Jiangsu province “Innovative and Entrepreneurial talent” and Six Talent Peaks Project of Jiangsu Province, the Emergency Prevention and Control Capacity Program for New Severe Infectious diseases of National Institute for Viral Disease Control and Prevention, and the 135 Strategic Program of Chinese Academy of Sciences, the Science and Technology Innovation Committee of Shenzhen Municipality (JCYJ20180228162229889).

## Author contributions

XW conducted most experiments, analyzed the data and wrote the draft manuscript. LC conducted all the neutralization experiments. BH, LZ, SX, HS, DZ, HY, WN provided technical assistance and did animal experiments. MF, YL, PY and QH evaluated the efficacy of Nbs in SARS-CoV-2 infected transgenic hACE2 mice. ZW designed the study, directed and financially supported the study and revised the manuscript. All authors critically reviewed the draft manuscript and approved the final version.

## Supplemental Materials

Materials and Methods

**Supplemental Figure 1. Characterization of anti-sera specific for SARS-CoV-2**. (**A**) The experimental schedule for immunization. The titer of anti-sera specific for SARS-CoV-2 S protein (**B**) and RBD protein (**C**) was evaluated one week after the immunization in alpaca receiving SARS-CoV-2 spike protein, respectively. The titer of the third anti-serum was indicated as blue line. The blue # indicates the anti-serum titer after the third immunization. 3^rd^ anti-serum and 2^nd^ anti-serum represent the anti-sera collected from alpaca one week after the 3^rd^ and 2^nd^ immunization. Blank serum represents the alpaca serum collected before immunization, which was taken as a negative control. (**D**) Neutralization potency of the immunized alpaca’s serum against pseudotyped SARS-CoV-2 was detected. ND_50_: half-maximal serum neutralization dilution titer. Titer and ND_50_ were indicated. Data of B-D represent as mean ± SEM. All experiments of B-D were repeated twice.

**Supplemental Figure 2. The construction and biopanning of C9-Nb library**. (**A**) The table summary of C9-Nb library, wherein phage displayed Nb of PBMC from alpaca receiving three times immunization of SARS-CoV-2 S protein. (**B**) The binding of the phage library with S via phage ELISA. Lib is the phage library of C9-Nb; 1^st^, 2^nd^, and 3^rd^ are the phage library after panning on 1 round, 2 rounds, and 3 rounds of S protein enrichment, respectively. (**C**) Single clone of phages from the C9-Nb library after the second and third enrichment of SARS-CoV-2 S were analyzed by phage ELISA. One dot represents the supernatant binding of one clone. Positive rate was indicated.

**Supplemental Figure 3. Characterization of Nb-Fc**. (**A**) The diagram of C9Nb, constituted by Nb fusing with human Fc1. (**B**) 21 various Nb-Fcs binding with S and RBD protein identified by ELISA. Grey dot represents negative control. Green dots represent the specific binding with S protein. Blue dots represent the double binding with S and RBD protein. (C) Representative binding curve of Nb-Fcs with RBD tested by BLI. (**D**) The table summary of 21 Nb-Fcs binding with RBD tested by BLI. (**E**) The cell supernatants of 21 various Nb-Fcs were tested for neutralization against SARS-CoV-2 infection, the cell supernatant displaying outstanding neutralizing curve was labeled as the color-coded curve. Data of B represent as mean ± SEM. All experiments of B-E were repeated twice

**Supplemental Figure 4. Characterization of purified Nb-Fcs**. (**A**) Purified Nb-Fcs binding with RBD identified by ELISA. Data represent as mean ± SEM. (**B**) RBD protein under reducing condition (R) or non-reducing condition (NR) was detected by WB with Nb_15_-Fc, Nb_22_-Fc and Nb_31_-Fc. Kinetic binding curve of RBD with Nb_15_-Fc (**C**), Nb_22_-Fc (**D**) and Nb_31_-Fc (**E**), respectively. Binding curves are colored black, and fit of the data to a 1:1 binding model is colored red.

**Supplemental Figure 5. Epitope analysis of Nb-Fcs by BLI**. RBD protein was coated on the sensor, Nb_15_-Fc (**A**), Nb_22_-Fc (**B**) or Nb_31_-Fc(C) as the first antibody was added to bind for 300 s, followed by the addition of Nb_15_-FC, Nb_22_-FC and Nb_31_-FC as the second antibody for another 300 s.

**Supplemental Figure S6. Characterizing the potency of neutralization against authentic SARS-CoV-2 conferred by Nb-Fcs**. The neutralization potency of Nb_15_-Fc (**A**), Nb_22_-Fc (**B**), Nb_31_-Fc (**C**), SNB02 (isotype control antibody) (**D**) was detected based on authentic SARS-CoV-2 plaque reduction neutralization test. The raw data was depicted. (**E**) A table summary authentic SARS-CoV-2 neutralization potencies of Nb-Fcs.

**Supplemental Figure 7. Characterization of Nb_15_s with multivalent or various formats**. (**A**) The binding curve of multivalent Nb_15_s with RBD protein detected by BLI. (**B**) The table summary of the binding of Nb_15_s with RBD protein tested by BLI. (**C**) Multivalent Nb_15_s and various formats were evaluated for neutralization potency against pseudotyped SARS-CoV-2 infection.

**Supplemental Figure 8. Kinetic binding curve of Nb_15_-Nb_H_-Nb_15_ with MSA**. Kinetic binding curve of Nb_15_-Nb_H_-Nb_15_ at the concentration of 300 nM, 100nM, 33.3 nM,11.1nM, 3.7nM and 1.2 nM with MSA by BLI. Binding curves are colored black, and fit of the data to a 1:1 binding model is colored red.

**Supplemental Table 1**. Summary of CDR sequences of positive Nb clones.

**Supplemental Table 2**. Summary of Nbs inhibiting SARS-CoV-2 variants.

**Supplemental Table 3**. Summary of various Nbs inhibiting pseudotyped SARS-CoV-2.

**Supplemental Table 4**. Summary of RBD binding with Nb_15_s in different conditions.

