## supplemental information for "A potent bispecific nanobody protects hACE2 mice against SARS-CoV-2 infection via intranasal administration"

### Supplemental Materials

Materials and Methods

Fig.S1

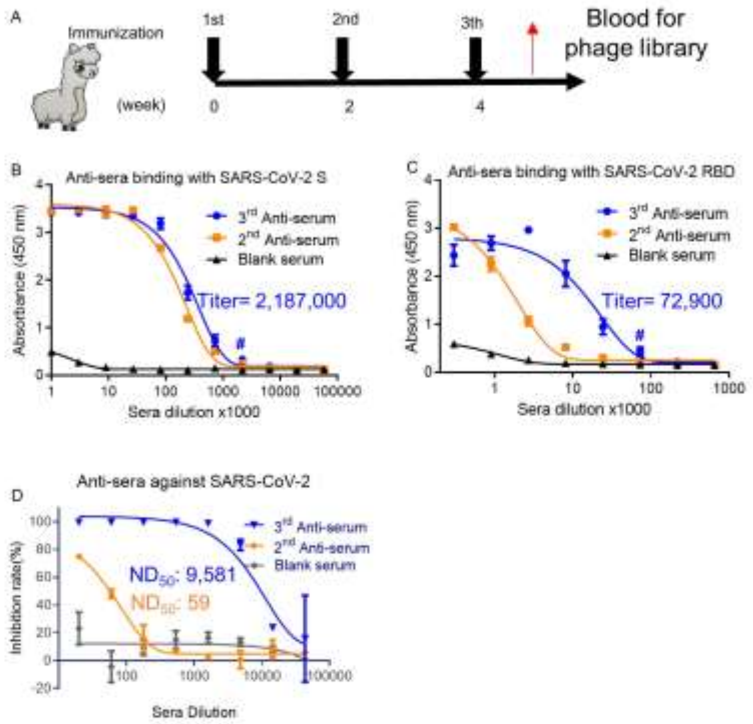


**Supplemental Figure 1. Characterization of anti-sera specific for SARS-CoV-2**. (**A**) The experimental schedule for immunization. The titer of anti-sera specific for SARS-CoV-2 S protein (**B**) and RBD protein (**C**) was evaluated one week after the immunization in alpaca receiving SARS-CoV-2 spike protein, respectively. The titer of the third anti-serum was indicated as blue line. The blue # indicates the anti-serum titer after the third immunization. 3$^{rd}$ anti-serum and 2$^{nd}$ anti-serum represent the anti-sera collected from alpaca one week after the 3$^{rd}$ and 2$^{nd}$ immunization. Blank serum represents the alpaca serum collected before immunization, which was taken as a negative control. (**D**) Neutralization potency of the immunized alpaca's serum against pseudotyped SARS-CoV-2 was detected. $ND_{50}$: half-maximal serum neutralization dilution titer. Titer and $ND_{50}$ were indicated. Data of B-D represent as mean $\pm$ SEM. All experiments of B-D were repeated twice.

Fig. S2

A

The summary of C9-Nb library

| Name | Library size | Sequencing clone | Sequence results | In frame clone | Diversity | In frame rate |
|---|---|---|---|---|---|---|
| C9-Nb-lib | 2.0*10^9 | 25 | 25 | 24 | 24/24 100% | 24/25 96% |

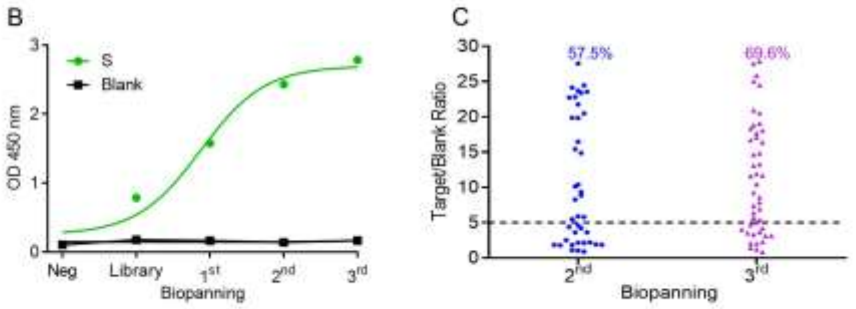


**Supplemental Figure 2. The construction and biopanning of C9-Nb library.** (**A**) The table summary of C9-Nb library, wherein phage displayed Nb of PBMC from alpaca receiving three times immunization of SARS-CoV-2 S protein. (**B**) The binding of the phage library with S via phage ELISA. Lib is the phage library of C9-Nb; 1$^{st}$, 2$^{nd}$, and 3$^{rd}$ are the phage library after panning on 1 round, 2 rounds, and 3 rounds of S protein enrichment, respectively. (**C**) Single clone of phages from the C9-Nb library after the second and third enrichment of SARS-CoV-2 S were analyzed by phage ELISA. One dot represents the supernatant binding of one clone. Positive rate was indicated.

### Fig. S3

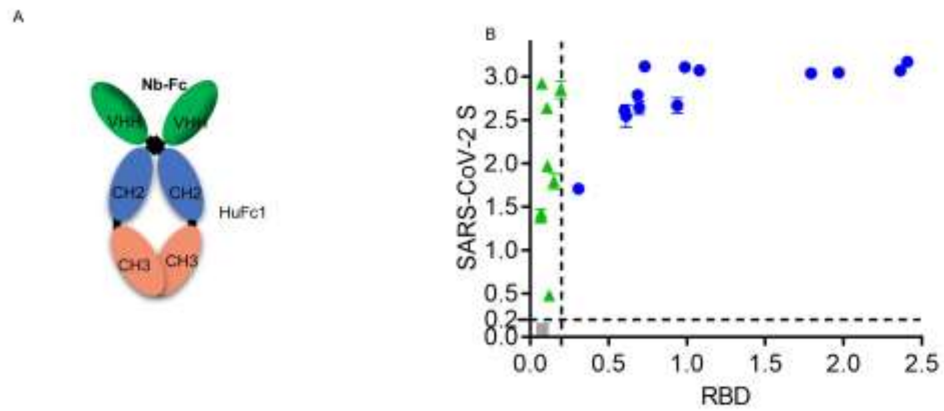


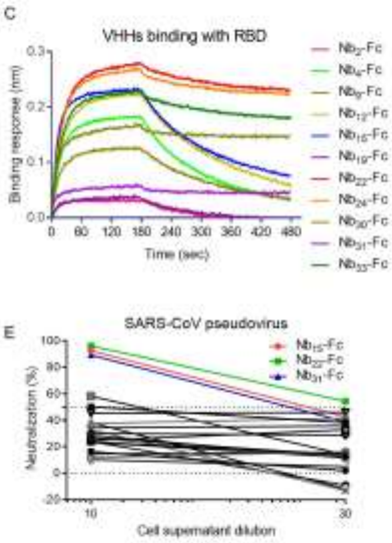


D

| Clone ID | Sample ID | Loading Sample ID | Response (nm) | KD (M) | kon(1/Ms) | kdis(1/s) | RMax | Full R^2 |
|---|---|---|---|---|---|---|---|---|
| $Nb_{1}$-Fc | RBD | S-1-1 | 0.0156 | 4.59E-08 | 6.48E+04 | 2.97E-03 | 0.0274 | 0.862 |
| $Nb_{2}$-Fc | RBD | S-2-1 | 0.0302 | 2.47E-08 | 6.85E+05 | 1.69E-02 | 0.0389 | 0.9273 |
| $Nb_{4}$-Fc | RBD | S-4-1 | 0.1824 | 1.43E-08 | 4.40E+05 | 6.30E-03 | 0.2024 | 0.9956 |
| $Nb_{5}$-Fc | RBD | S-5-1 | 0.0195 | 1.46E-09 | 7.62E+05 | 1.11E-03 | 0.0176 | 0.3834 |
| $Nb_{7}$-Fc | RBD | S-7-1 | 0.0166 | 1.30E-07 | 1.84E+05 | 2.40E-02 | 0.0432 | 0.7633 |
| $Nb_{9}$-Fc | RBD | S-9-1 | 0.1254 | 9.04E-09 | 5.16E+05 | 4.67E-03 | 0.1327 | 0.9938 |
| $Nb_{12}$-Fc | RBD | S-12-1 | 0.2218 | 1.24E-08 | 3.79E+05 | 4.70E-03 | 0.2443 | 0.9966 |
| $Nb_{13}$-Fc | RBD | S-13-2 | 0.0203 | 8.19E-10 | 3.04E+05 | 2.49E-04 | 0.0192 | 0.5595 |
| $Nb_{14}$-Fc | RBD | S-14-1 | 0.0159 | <1.0E-12 | 5.76E+04 | <1.0E-07 | 0.0271 | 0.8809 |
| $Nb_{15}$-Fc | RBD | S-15-1 | 0.2314 | 6.09E-09 | 6.94E+05 | 4.22E-03 | 0.2366 | 0.9879 |
| $Nb_{19}$-Fc | RBD | S-19-1 | 0.0366 | 3.76E-08 | 3.48E+05 | 1.31E-02 | 0.0493 | 0.9862 |
| $Nb_{22}$-Fc | RBD | S-22-1 | 0.2322 | 5.99E-09 | 4.92E+05 | 2.94E-03 | 0.2367 | 0.9927 |
| $Nb_{24}$-Fc | RBD | S-24-1 | 0.1684 | 4.25E-09 | 8.30E+05 | 3.53E-03 | 0.1706 | 0.9876 |
| $Nb_{25}$-Fc | RBD | S-25-1 | 0.0168 | 6.14E-09 | 3.12E+05 | 1.92E-03 | 0.0169 | 0.5564 |
| $Nb_{27}$-Fc | RBD | S-27-1 | 0.016 | 2.28E-08 | 2.46E+05 | 5.61E-03 | 0.0199 | 0.8945 |
| $Nb_{28}$-Fc | RBD | S-28-1 | 0.0068 | 2.86E-05 | 2.04E+03 | 5.82E-02 | 1.7237 | 0.3166 |
| $Nb_{29}$-Fc | RBD | S-29-1 | 0.0169 | 4.45E-09 | 1.34E+05 | 5.95E-04 | 0.0204 | 0.8342 |
| $Nb_{30}$-Fc | RBD | S-30-1 | 0.0469 | 2.12E-08 | 5.46E+05 | 1.16E-02 | 0.0525 | 0.9855 |
| $Nb_{31}$-Fc | RBD | S-31-1 | 0.246 | 1.04E-08 | 6.37E+05 | 6.64E-03 | 0.261 | 0.9823 |
| $Nb_{33}$-Fc | RBD | S-33-1 | 0.1328 | 4.51E-09 | 4.34E+05 | 1.95E-03 | 0.1364 | 0.994 |
| $Nb_{34}$-Fc | RBD | S-34-2 | 0.0114 | 8.46E-06 | 1.73E+03 | 1.47E-02 | 0.975 | 0.8974 |

**Supplemental Figure 3. Characterization of Nb-Fc.** (**A**) The diagram of C9Nb, constituted by Nb fusing with human Fc1. (**B**) 21 various Nb-Fcs binding with S and RBD protein identified by ELISA. Grey dot represents negative control. Green dots represent the specific binding with S protein. Blue dots represent the double binding with S and RBD protein. (C) Representative binding curve of Nb-Fcs with RBD tested by BLI. (**D**) The table summary of 21 Nb-Fcs binding with RBD tested by BLI. (**E**) The cell supernatants of 21 various Nb-Fcs were tested for neutralization against SARS-CoV-2 infection, the cell supernatant displaying outstanding neutralizing curve was labeled as the color-coded curve. Data of B represent as mean ± SEM. All experiments of B-E were repeated twice

Fig.S4

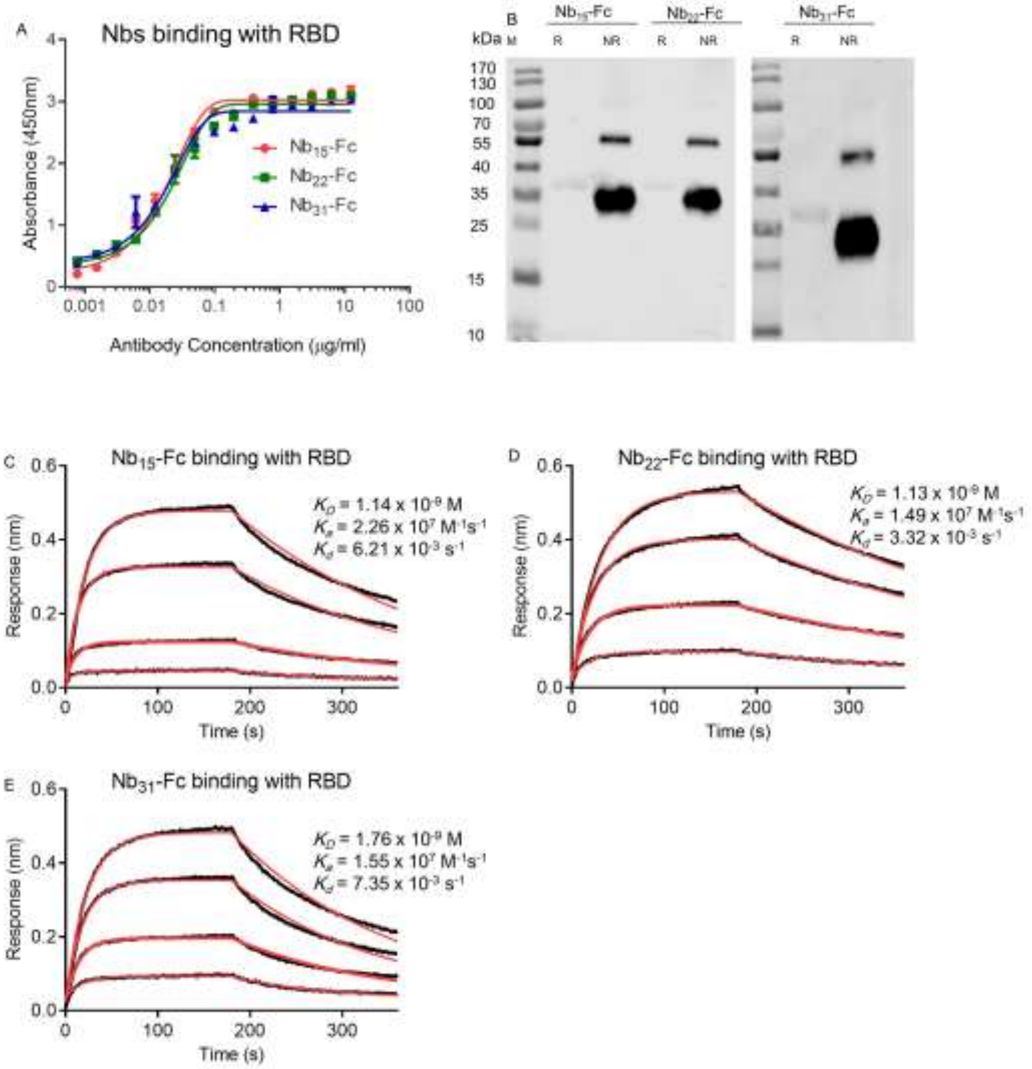


**Supplemental Figure 4. Characterization of purified Nb-Fcs.** (**A**) Purified Nb-Fcs binding with RBD identified by ELISA. Data represent as mean ± SEM. (**B**) RBD protein under reducing condition (R) or non-reducing condition (NR) was detected by WB with $Nb_{15}$-Fc, $Nb_{22}$-Fc and $Nb_{31}$-Fc. Kinetic binding curve of RBD with $Nb_{15}$-Fc (**C**), $Nb_{22}$-Fc (**D**) and $Nb_{31}$-Fc (**E**), respectively. Binding curves are colored black, and fit of the data to a 1:1 binding model is colored red.

Fig. S5

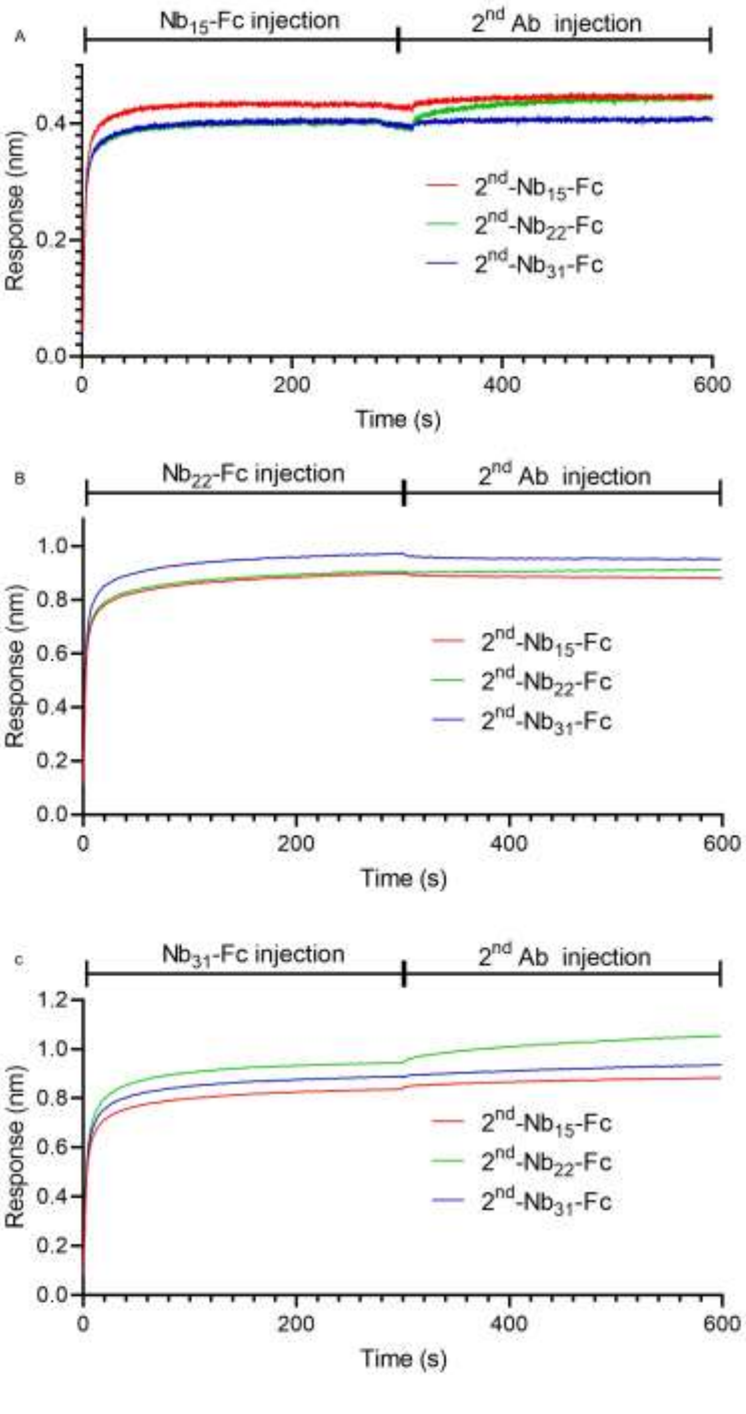


**Supplemental Figure 5. Epitope analysis of Nb-Fcs by BLI.** RBD protein was coated on the sensor, $Nb_{15}$-Fc (**A**), $Nb_{22}$-Fc (**B**) or $Nb_{31}$-Fc(C) as the first antibody was added to bind for 300 s, followed by the addition of $Nb_{15}$-FC, $Nb_{22}$-FC and $Nb_{31}$-FC as the second antibody for another 300 s.

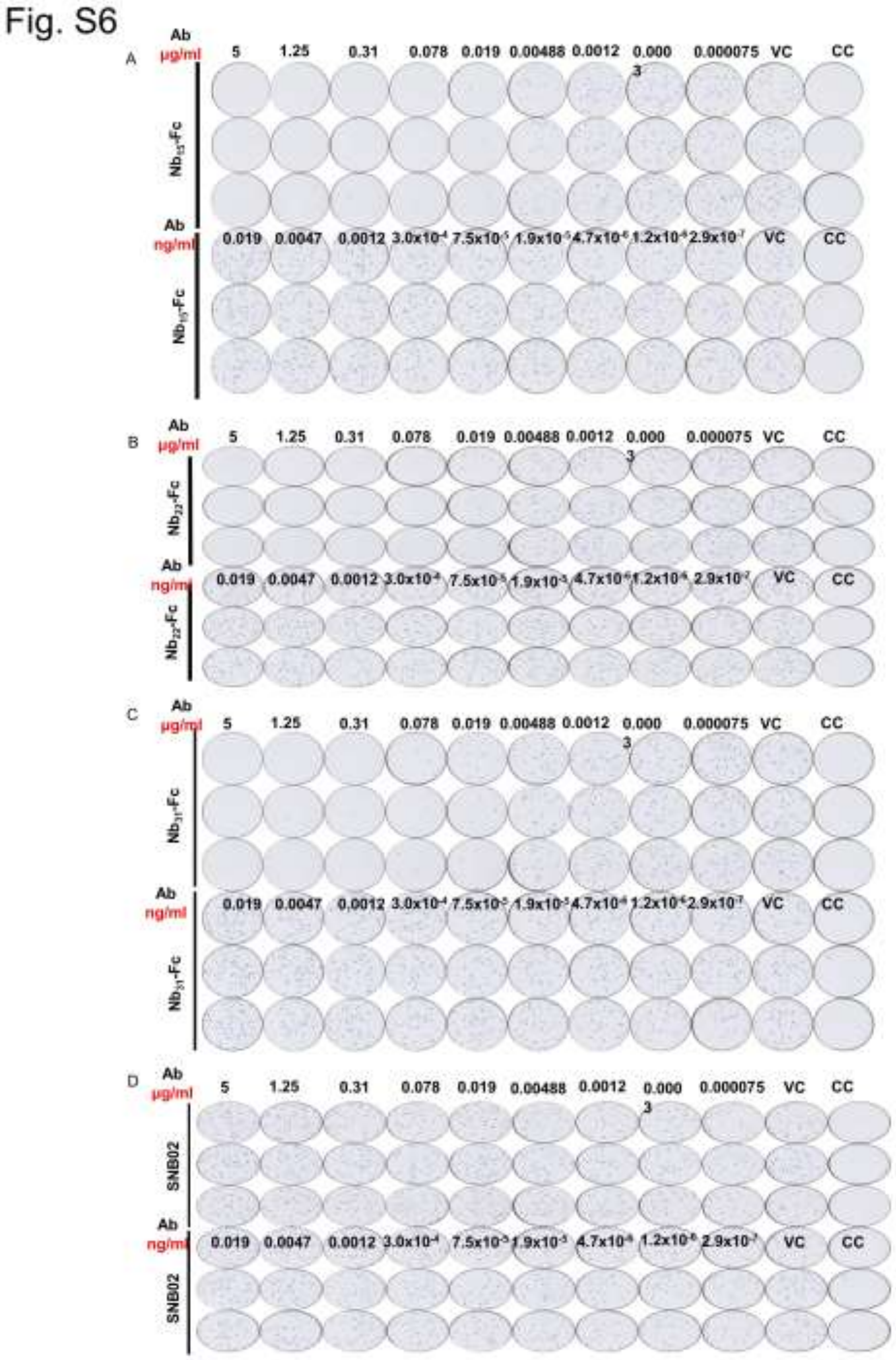


E Summary of C9Nbs inhibiting SARS-CoV-2 live virus

| μg/ml | SARS-CoV-2 live virus | | |
|---|---|---|---|
| | $IC_{50}$ | $IC_{80}$ | $IC_{90}$ |
| $Nb_{15}$-Fc | 0.0033 | 0.0084 | 0.0156 |
| $Nb_{22}$-Fc | 0.0068 | 0.0108 | 0.0159 |
| $Nb_{31}$-Fc | 0.0036 | 0.0100 | 0.0235 |

**Supplemental Figure S6. Characterizing the potency of neutralization against authentic SARS-CoV-2 conferred by Nb-Fcs.** The neutralization potency of $Nb_{15}$-Fc (**A**), $Nb_{22}$-Fc (**B**), $Nb_{31}$-Fc(**C**), SNB02 (isotype control antibody) (**D**) was detected based on authentic SARS-CoV-2 plaque reduction neutralization test. The raw data was depicted. (**E**) A table summary authentic SARS-CoV-2 neutralization potencies of Nb-Fcs.

Fig.S7

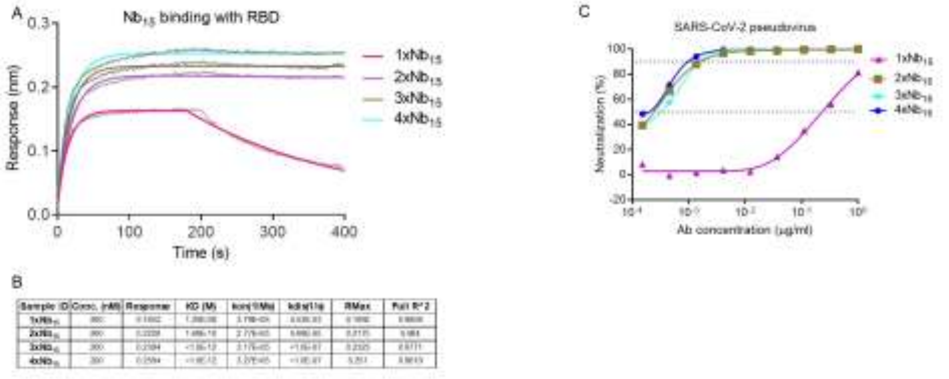


**Supplemental Figure 7. Characterization of $Nb_{15}$s with multivalent or various formats.** (**A**) The binding curve of multivalent $Nb_{15}$s with RBD protein detected by BLI. (**B**) The table summary of the binding of $Nb_{15}$s with RBD protein tested by BLI. **(C)** Multivalent $Nb_{15}$s and various formats were evaluated for neutralization potency against pseudotyped SARS-CoV-2 infection.

Fig. S8

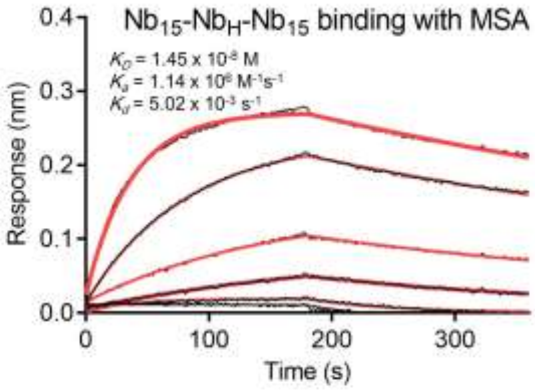


**Supplemental Figure 8. Kinetic binding curve of $Nb_{15}$-$Nb_H$-$Nb_{15}$ with MSA.** Kinetic binding curve of $Nb_{15}$-$Nb_H$-$Nb_{15}$ at the concentration of 300 nM, 100nM, 33.3 nM,11.1nM, 3.7nM and 1.2 nM with MSA by BLI. Binding curves are colored black, and fit of the data to a 1:1 binding model is colored red.

**Supplemental Table 1**. Summary of CDR sequences of positive Nb clones.

| ID | CDR1 | CDR2 | CDR3 |
|---|---|---|---|
| $Nb_1$-Fc | GNIFSIYT | VTSGGST | N—ARLFDPGY |
| $Nb_2$-Fc | GGTLASFA | INIINRT | AAHFVPPGSRLRDCLVNELYNY |
| $Nb_4$-Fc | GGTLASFA | INIINRT | AAHFVPPGSRLRGCLVNELYNY |
| Nb5-Fc | GFTWNYHA | ISSSGSTT | AAPHSGSVCPR-WAEYYGVDH |
| $Nb_7$-Fc | GGTLASFA | INIINRT | AAHFVPPGSRLRGCLVNEAYNY |
| Nb9-Fc | GGTLASFA | INIINRT | AAHFVPPGGRLRGCLVNDLYNY |
| $Nb_{12}$-Fc | GGTLASFA | INIINRT | AAHFVPPGSRLRGCLVNDLYNY |
| $Nb_{13}$-Fc | KIILSFYD | ITNSGST | N—TFHY |
| $Nb_{14}$-Fc | GFTSDRYT | ISSSGGST | TARPSLWAVVAGCPLDQNTYFS |
| $Nb_{15}$-Fc | GFTLDYYA | ISSSGST | AG-VVHDVQAM-CVMNP-WGS |
| $Nb_{19}$-Fc | GGTLASFA | INIINRT | AAHFVPPGSRLRGCLVNDVYNY |
| $Nb_{22}$-Fc | GGTLASFA | IDVINRA | AAHFVPPGSRLRGCLVNELYNY |
| $Nb_{24}$-Fc | GGTLASFA | INIINRT | AAHFVPPESRLRGCLVNELYNY |
| $Nb_{25}$-Fc | FSISSIDT | ITSRRDT | YG—QDVLGQIY |
| $Nb_{27}$-Fc | RNIFSIYT | ITSGGST | TT—AGSWQGDY |
| $Nb_{28}$-Fc | GGTLASFA | INIINRT | AAHFVPPESRLRGCLVNEAYNY |
| $Nb_{29}$-Fc | FSISSVDT | ISSRSFT | YG—QDILGQIY |
| $Nb_{30}$-Fc | GTLASFA | INIINRT | AAHFVPPGSRLRGCLVNELYNY |
| $Nb_{31}$-Fc | GGTLASFA | INIINRP | AAHFVPPGSRLGGCLVNELYNY |
| $Nb_{33}$-Fc | GGTLASFA | INIINRT | AAHFVPPGSRFRGCSVNELYNY |
| $Nb_{34}$-Fc | GLTLHLYD | INIINRP | AAHFVPPGSRLGGCLVNELYNY |

**Supplemental Table 2**. Summary of Nbs inhibiting SARS-CoV-2 variants.

| Variants | $Nb_{15}$-Fc (mean±sd μg/ml) | | | $Nb_{22}$-Fc (mean±sd μg/ml) | | | $Nb_{31}$-Fc (mean±sd μg/ml) | | | $Nb_{15}$-$Nb_H$-$Nb_{15}$ (mean±sd μg/ml) | | |
|---|---|---|---|---|---|---|---|---|---|---|---|---|
| | $IC_{50}$ | $IC_{80}$ | $IC_{90}$ | $IC_{50}$ | $IC_{80}$ | $IC_{90}$ | $IC_{50}$ | $IC_{80}$ | $IC_{90}$ | $IC_{50}$ | $IC_{80}$ | $IC_{90}$ |
| WT | 0.0008±0.0001 | 0.0019±0.0004 | 0.0033±0.0012 | 0.0016±0.0001 | 0.0046±0.0012 | 0.0086±0.0033 | 0.0023±0.0004 | 0.0083±0.0019 | 0.0183±0.0059 | 0.0004±0 | 0.0012±0.0004 | 0.0018±0.0009 |
| Q321L | 0.0009±0.0004 | 0.0023±0.0007 | 0.0039±0.0008 | 0.0014±0.0003 | 0.0042±0.0009 | 0.0079±0.0019 | 0.002±0.0005 | 0.0065±0.0017 | 0.0133±0.0033 | 0.001±0.0001 | 0.0022±0.0004 | 0.0034±0.0009 |
| V341I | 0.0007±0.0002 | 0.0026±0.0005 | 0.0059±0.0019 | 0.0017±0.0005 | 0.0042±0.0007 | 0.007±0.001 | 0.0028±0.0004 | 0.0087±0.0024 | 0.0169±0.0058 | 0.0011±0.0001 | 0.0027±0.0009 | 0.0047±0.0022 |
| A348T | 0.001±0.0002 | 0.0023±0.0003 | 0.0036±0.0005 | 0.0019±0.0008 | 0.0046±0.0012 | 0.0076±0.0016 | 0.0029±0.0001 | 0.0088±0.0013 | 0.0176±0.0032 | 0.0008±0.0002 | 0.0014±0.0001 | 0.0019±0.0004 |
| N354D | 0.0008±0.0002 | 0.0022±0.0002 | 0.0041±0.001 | 0.0013±0.0003 | 0.0033±0.0005 | 0.0056±0.0012 | 0.0019±0.0001 | 0.0068±0.0018 | 0.0145±0.0051 | 0.0006±0.0002 | 0.0016±0.0006 | 0.0032±0.0013 |
| S359N | 0.0011±0.0001 | 0.0026±0.0003 | 0.0043±0.0006 | 0.0016±0.0003 | 0.0037±0.0005 | 0.0061±0.0008 | 0.002±0.0005 | 0.0064±0.0012 | 0.0129±0.0037 | 0.0008±0.0002 | 0.0017±0.0002 | 0.0025±0.0005 |
| V367F | 0.0007±0.0001 | 0.002±0.0002 | 0.0036±0.0004 | 0.0011±0.0003 | 0.0033±0.0003 | 0.0062±0.0004 | 0.0021±0.0006 | 0.0095±0.0017 | 0.0237±0.0022 | 0.0005±0 | 0.0013±0.0001 | 0.0026±0.0006 |
| K378R | 0.0007±0.0002 | 0.0024±0.0003 | 0.0046±0.0001 | 0.0012±0.0002 | 0.0028±0.0003 | 0.0045±0.0005 | 0.0013±0.0001 | 0.004±0.0004 | 0.0079±0.0017 | 0.0008±0.0002 | 0.002±0.0004 | 0.0033±0.0006 |
| R408I | 0.0007±0.0002 | 0.002±0.0007 | 0.0035±0.0016 | 0.001±0.0001 | 0.0027±0.0003 | 0.0047±0.0009 | 0.0014±0.0002 | 0.0038±0.001 | 0.0069±0.0026 | 0.0007±0.0001 | 0.0014±0.0002 | 0.002±0.0005 |
| Q409E | 0.0005±0.0001 | 0.0014±0 | 0.0026±0.0002 | 0.0009±0.0003 | 0.0022±0.0007 | 0.0036±0.0011 | 0.0009±0.0001 | 0.0029±0.0004 | 0.0057±0.0007 | 0.0007±0.0002 | 0.0014±0.0002 | 0.0021±0.0003 |
| K458R | 0.0009±0.0003 | 0.0025±0.0002 | 0.0047±0.0006 | 0.0013±0.0001 | 0.0041±0.0014 | 0.008±0.0036 | 0.0025±0.0003 | 0.0068±0.0013 | 0.0128±0.004 | 0.0008±0.0002 | 0.0018±0.0003 | 0.0029±0.0004 |
| G476S | 0.0006±0.0002 | 0.0015±0.0004 | 0.0025±0.0006 | 0.0013±0.0006 | 0.0034±0.0011 | 0.006±0.0013 | 0.0015±0.0001 | 0.0049±0.0008 | 0.0101±0.0023 | 0.0003±0.0001 | 0.0009±0.0002 | 0.0014±0.0003 |
| V483A | 0.0006±0.0001 | 0.0017±0.0005 | 0.0031±0.0011 | 0.0015±0.0003 | 0.0039±0.0008 | 0.0069±0.0026 | 0.0027±0.001 | 0.0093±0.0018 | 0.0206±0.0027 | 0.0005±0.0001 | 0.0013±0 | 0.0022±0.0002 |
| Y508H | 0.0005±0.0001 | 0.0023±0.0003 | 0.0053±0.001 | 0.0013±0.0002 | 0.0035±0.0007 | 0.006±0.0015 | 0.0018±0.0005 | 0.0068±0.0007 | 0.0157±0.0047 | 0.0008±0.0001 | 0.0019±0.0006 | 0.0033±0.0016 |
| H519P | 0.0008±0.0001 | 0.0023±0.0002 | 0.0041±0.0005 | 0.0011±0.0002 | 0.0033±0 | 0.006±0.0004 | 0.0014±0.0003 | 0.005±0.0004 | 0.0107±0.0032 | 0.0007±0.0002 | 0.0016±0.0001 | 0.0025±0.0002 |
| D614G | 0.0007±0.0001 | 0.002±0.0002 | 0.0035±0.0007 | 0.0012±0.0005 | 0.0033±0.0005 | 0.0059±0.0003 | 0.0015±0.0004 | 0.0039±0.001 | 0.0067±0.0016 | 0.0006±0.0001 | 0.0015±0.0005 | 0.0026±0.001 |
| A435S | N/A | N/A | N/A | N/A | N/A | N/A | N/A | N/A | N/A | 0.0007±0.0001 | 0.0015±0.0003 | 0.0024±0.0006 |
| I472V | N/A | N/A | N/A | N/A | N/A | N/A | N/A | N/A | N/A | 0.0006±0.0001 | 0.0014±0.0003 | 0.0023±0.0006 |
| N501Y | N/A | N/A | N/A | N/A | N/A | N/A | N/A | N/A | N/A | 0.0007±0.0002 | 0.002±0.0005 | 0.0042±0.0013 |

*Note: N/A, no test.*

**Supplemental Table 3**. Summary of various Nbs inhibiting pseudotyped SARS-CoV-2.

| Nbs | $IC_{50}$ (mean±sd) | $IC_{80}$ (mean±sd) | $IC_{90}$ (mean±sd) |
|---|---|---|---|
| $1xNb_{15}$ | 0.3074±0.0237 | 0.5059±0.0699 | 0.622±0.0969 |
| $2xNb_{15}$ | 0.0003±0 | 0.0008±0.0001 | 0.0011±0.0001 |
| $3xNb_{15}$ | 0.0004±0 | 0.001±0 | 0.0014±0.0001 |
| $4xNb_{15}$ | 0.0002±0.0001 | 0.0008±0.0001 | 0.0011±0.0002 |
| $Nb_{15}$-$Nb_H$ | 0.5529±0.0889 | 1.071±0.1754 | 1.374±0.2263 |
| $Nb_H$-$Nb_{15}$ | 0.1974±0.004 | 0.3469±0.0533 | 0.4344±0.0822 |
| $Nb_{15}$-$Nb_{15}$-$Nb_H$ | 0.0251±0.0058 | 0.0419±0.0051 | 0.0517±0.0047 |
| $Nb_H$-$Nb_{15}$-$Nb_{15}$ | 0.0008±0.0001 | 0.0017±0.0003 | 0.0022±0.0004 |
| $Nb_{15}$-$Nb_H$-$Nb_{15}$ | 0.0004±0 | 0.0009±0 | 0.0013±0 |
| $Nb_{15}$-Fc | 0.0009±0.0001 | 0.0019±0.0001 | 0.0025±0.0002 |

μg/ml

<0.001

0.001-0.01

0.01-0.1

>0.1

**Supplemental Table 4**. Summary of RBD binding with $Nb_{15}$s in different conditions.

| Sample ID | Condition | Conc. (nM) | Response | KD (M) | kon (1/Ms) | kdis (1/s) | RMax | Full R^2 |
|---|---|---|---|---|---|---|---|---|
| $Nb_{15}$-$Nb_H$-$Nb_{15}$ | WT | 133.3 | 0.3806 | <1.0E-12 | 3.44E+05 | <1.0E-07 | 0.3687 | 0.9676 |
| | Aero | 133.3 | 0.2139 | 9.92E-09 | 4.11E+04 | 4.08E-04 | 0.3277 | 0.9957 |
| | 37 °C | 133.3 | 0.3756 | <1.0E-12 | 1.89E+05 | <1.0E-07 | 0.374 | 0.9948 |
| | 50 °C | 133.3 | 0.3903 | 5.78E-11 | 2.11E+05 | 1.22E-05 | 0.3849 | 0.996 |
| | 60 °C | 133.3 | 0.4048 | 5.57E-10 | 1.91E+05 | 1.06E-04 | 0.402 | 0.9986 |
| | 70 °C | 133.3 | 0.3775 | 2.52E-10 | 1.63E+05 | 4.10E-05 | 0.3801 | 0.9989 |
| | 80 °C | 133.3 | 0.3199 | 5.64E-09 | 5.85E+04 | 3.30E-04 | 0.4193 | 0.9994 |
| | 90 °C | 133.3 | 0.1078 | 1.02E-08 | 2.90E+04 | 2.95E-04 | 0.211 | 0.9978 |
| $Nb_{15}$-Fc | WT | 62.5 | 0.8028 | <1.0E-12 | 6.98E+05 | <1.0E-07 | 0.7814 | 0.9824 |
| | Aero | 62.5 | 0.3398 | 2.79E-10 | 5.05E+04 | 1.41E-05 | 0.7387 | 0.999 |
| | 37 °C | 62.5 | 0.8136 | <1.0E-12 | 4.34E+05 | <1.0E-07 | 0.804 | 0.9946 |
| | 50 °C | 62.5 | 0.853 | <1.0E-12 | 4.11E+05 | <1.0E-07 | 0.848 | 0.9976 |
| | 60 °C | 62.5 | 0.7989 | 4.38E-11 | 4.11E+05 | 1.80E-05 | 0.7944 | 0.9979 |
| | 70 °C | 62.5 | 0.6126 | 4.95E-09 | 8.87E+04 | 4.39E-04 | 0.956 | 0.9995 |
| | 80 °C | 62.5 | 0.2077 | 1.98E-08 | 4.16E+04 | 8.26E-04 | 0.557 | 0.9983 |
| | 90 °C | 62.5 | 0.1088 | 4.12E-08 | 2.76E+04 | 1.14E-03 | 0.4281 | 0.9978 |
| $3xNb_{15}$ | WT | 133.3 | 0.3769 | <1.0E-12 | 6.20E+05 | <1.0E-07 | 0.3637 | 0.8833 |
| | Aero | 133.3 | 0.184 | <1.0E-12 | 2.55E+04 | <1.0E-07 | 0.3816 | 0.9946 |
| | 37 °C | 133.3 | 0.3667 | <1.0E-12 | 4.01E+05 | <1.0E-07 | 0.3578 | 0.9563 |
| | 50 °C | 133.3 | 0.3588 | <1.0E-12 | 3.96E+05 | <1.0E-07 | 0.3507 | 0.9583 |
| | 60 °C | 133.3 | 0.3715 | <1.0E-12 | 3.49E+05 | <1.0E-07 | 0.3635 | 0.9717 |
| | 70 °C | 133.3 | 0.2294 | <1.0E-12 | 2.78E+04 | <1.0E-07 | 0.4508 | 0.9975 |
| | 80 °C | 133.3 | 0.2748 | 4.97E-10 | 4.53E+04 | 2.25E-05 | 0.4052 | 0.9994 |
| | 90 °C | 133.3 | 0.1294 | 4.87E-09 | 5.21E+04 | 2.54E-04 | 0.1778 | 0.9968 |
